# The evolutionary history of 2,658 cancers

**DOI:** 10.1101/161562

**Authors:** Moritz Gerstung, Clemency Jolly, Ignaty Leshchiner, Stefan C. Dentro, Santiago Gonzalez, Daniel Rosebrock, Thomas J. Mitchell, Yulia Rubanova, Pavana Anur, Kaixian Yu, Maxime Tarabichi, Amit Deshwar, Jeff Wintersinger, Kortine Kleinheinz, Ignacio Vázquez-García, Kerstin Haase, Lara Jerman, Subhajit Sengupta, Geoff Macintyre, Salem Malikic, Nilgun Donmez, Dimitri G. Livitz, Marek Cmero, Jonas Demeulemeester, Steven Schumacher, Yu Fan, Xiaotong Yao, Juhee Lee, Matthias Schlesner, Paul C. Boutros, David D. Bowtell, Hongtu Zhu, Gad Getz, Marcin Imielinski, Rameen Beroukhim, S. Cenk Sahinalp, Yuan Ji, Martin Peifer, Florian Markowetz, Ville Mustonen, Ke Yuan, Wenyi Wang, Quaid D. Morris, Paul T. Spellman, David C. Wedge, Peter Van Loo, on behalf of the PCAWG Evolution and Heterogeneity Working Group, the PCAWG network

## Abstract

Cancer develops through a process of somatic evolution. Here, we use whole-genome sequencing of 2,778 tumour samples from 2,658 donors to reconstruct the life history, evolution of mutational processes, and driver mutation sequences of 39 cancer types. The early phases of oncogenesis are driven by point mutations in a small set of driver genes, often including biallelic inactivation of tumour suppressors. Early oncogenesis is also characterised by specific copy number gains, such as trisomy 7 in glioblastoma or isochromosome 17q in medulloblastoma. By contrast, increased genomic instability, a nearly four-fold diversification of driver genes, and an acceleration of point mutation processes are features of later stages. Copy-number alterations often occur in mitotic crises leading to simultaneous gains of multiple chromosomal segments. Timing analysis suggests that driver mutations often precede diagnosis by many years, and in some cases decades, providing a window of opportunity for early cancer detection.

## Introduction

Akin to the evolution of species, the ∼10^15^ cells in the human body are subject to the forces of mutation and selection^1^. This process of somatic evolution begins before birth in the zygote and only comes to rest at death, as each cell division typically introduces between 1-10 mutations^2^. While these mutations are predominantly selectively neutral passenger mutations without transformative capacity, some are proliferatively advantageous driver mutations leading to cancer^3,4^. While the types of mutations in cancer genomes are well studied, little is known about the times when these lesions arise during somatic evolution and where the boundary between normal evolution and cancer progression should be drawn.

Sequencing of bulk tumour samples enables partial reconstruction of the evolutionary history of individual tumours, based on the catalogue of somatic mutations they have accumulated^5^. The timing of chromosomal gains during early somatic evolution and tumour progression can be estimated by counting point mutations within duplicated regions in single cancer samples^6-8^. Moreover, many studies have focused on the late stages of cancer evolution by reconstructing phylogenetic relationships between tumour samples and metastases from individual patients^9-12^. In addition, the typical ordering of driver mutations within a tumour type can be determined by aggregating pairwise timing estimates of genomic changes (for example clonal *vs*. subclonal) across many samples using preference models^13,14^.

Here, we extend and apply these methods to the Pan-Cancer Analysis of Whole Genomes (PCAWG)^15^ dataset, as part of the International Cancer Genome Consortium (ICGC)^16^ and The Cancer Genome Atlas (TCGA)^17^ to characterise the evolutionary history of 2,778 cancers from 2,658 unique donors across 39 cancer types (**Methods** and **Supplementary Information**). We determine the timing and order of mutations during cancer development to delineate patterns of chromosomal evolution within and across cancer types. We then define broad periods of tumour evolution and examine how drivers and mutational signatures vary between these stages. Finally, using clock-like mutational processes, we map mutation timing estimates into approximate real time, and create typical timelines of tumour evolution.

## Results

### Reconstructing the life history of single tumours

A cancer cell’s genome is shaped by the cumulative somatic aberrations that have arisen during its evolutionary past, and part of this history can be reconstructed from whole genome sequencing data^5^ (**Fig. 1a**). Initially, each point mutation occurs on a single chromosome in a single cell, which gives rise to a lineage of cells all bearing the same mutation. If that chromosomal locus is subsequently duplicated in this lineage, and the point mutation is on the allele that is gained, the mutation will also be duplicated, and both mutated copies can be detected in sequencing data. Mutations on other chromosomes, or on the non-duplicated allele of a gained region, will be present only as a single copy. Mutations found in a subset of tumour cells have not swept through the population and must have occurred after the most recent common ancestor (MRCA) of the tumour cells in the sequenced sample. Based on these principles, we are able to approximately time individual point mutations relative to other genomic events such as large-scale chromosomal gains, and the emergence of the MRCA.

**Figure 1.**
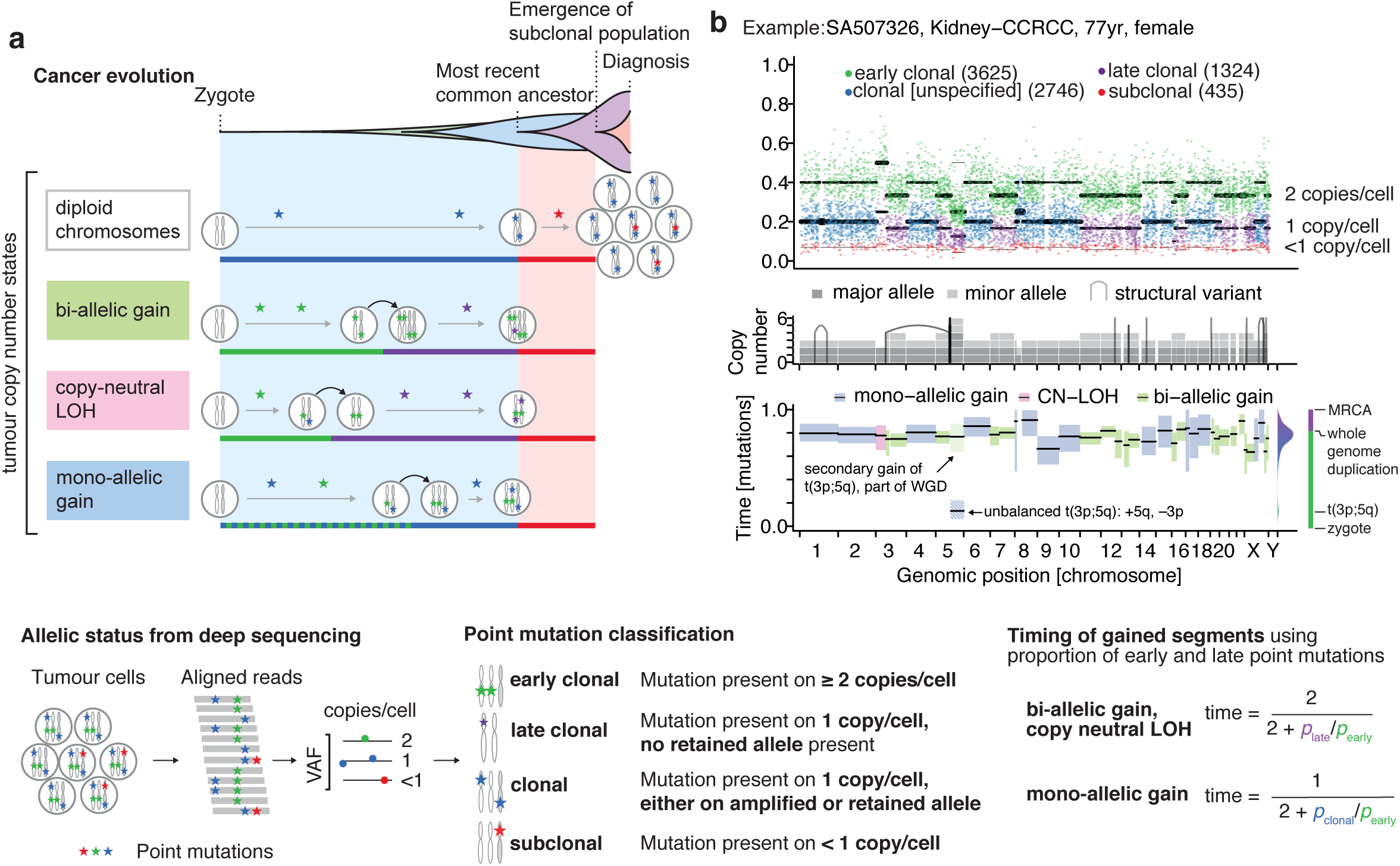
Principles of timing mutations. (**a**) Principles of timing mutations based on whole genome sequencing. According to the clonal evolution model of cancer, tumour cells evolve in multiple selective sweeps. During some of these sweeps, copy number gains are acquired, which can be used for timing analyses (green and purple epochs). Mutations acquired after the last clonal expansion are present in distinct subclonal populations (red epoch). The number of sequencing reads reporting point mutations can be used to discriminate variants as early or late clonal (green/purple) in cases of specific copy number gains, as well as clonal (blue) or subclonal (red) in cases without (right). The distribution of the number of early and late clonal mutations carries information about the timing of the copy number gains with the exact relation depending on the resulting copy number configuration (bottom). (**b**) Example case illustrating the annotation of point mutations based on the variant allele frequency (VAF, top) and copy number configuration including structural variants (middle). Each is shown as a function of genomic coordinate (x-axis). The resulting timing estimates for each copy number segment are shown at the bottom, indicating that, after an initial gain of chromosome arm 5q as part of an unbalanced translocation, a whole genome duplication occurred which gained all existing chromosomes in a single event.

Mapping point mutations to the proportion of cells and chromosomes enables us to define broad categories, each associated with broad epochs of tumour evolution (**Fig. 1a**). *Clonal* mutations have occurred before the emergence of the MRCA and are common to all cancer cells, whilst *subclonal* mutations are only observed in a fraction of cancer cells. In regions of clonal copy number gains, clonal mutations can be further classified as *early clonal* or *late clonal. Early clonal mutations* lie within a duplicated locus and are themselves doubled, *i.e.* they occurred prior to the copy number gain. *Late clonal mutations* also reside at a gained chromosomal region, but only as one copy, *i.e.* they happened after the copy number gain. Inevitably, there will be some crossover between these categories of clonal mutations, both within and between samples, as their timing depends on the timing of the surrounding copy number gains, which may take place at different times during tumour evolution. However, the broad comparison of mutations before and after copy number gains does give additional insight into the dynamics of clonal evolution.

Importantly, the relative frequency of early and late clonal mutations provides information about the timing of the underlying copy number segment. The ratio of doubled to non-doubled mutations within a gained region can be used to calculate a point estimate for when the gain happened during clonal evolution, referred to here as *mutational timing*. For example, there would be few, if any, co-amplified early clonal mutations if the gain had occurred right after fertilisation, whilst a gain that happened towards the end of clonal tumour evolution would contain many duplicated mutations^6^ (**Fig. 1a** and **Online Methods**). Later on, we will show how this mutational timing relates to real time for certain evolutionary events.

These analyses are illustrated in **Fig. 1b**. As expected, the variant allele frequency (VAF) of somatic point mutations cluster around the values imposed by the purity of the sample, local copy number configuration and identified subclonal population. The depicted clear cell renal cell carcinoma has gained chromosome arm 5q at an early time as part of an unbalanced translocation of t(3p;5q), confirming the notion that this lesion often occurs in adolescence in this cancer type^8^. At a later time point the sample underwent a whole genome duplication (WGD), duplicating all alleles including the derivative chromosome in a single event, as evidenced by the mutation time estimates of all copy number gains clustering around a single time point, independently of the exact copy number state.

### Timing patterns of copy number gains

To systematically explore the timing of copy number gains across cancer types, we applied mutational timing analysis to 2,111 samples from the PCAWG dataset that had copy number gains suitable for timing (see **Supplementary Information**). We find that chromosomal gains typically occur after more than half of the point mutations have been acquired (median value 0.76, inter-quartile range IQR [0.43, 0.94], with systematic differences between tumour types (**Fig. 2a**, **Extended Data Fig. 1**). In glioblastoma, medulloblastoma and pancreatic neuroendocrine cancers, a substantial fraction of gains occur early in mutational time. Conversely, in squamous cell lung cancers and melanomas, gains arise towards the end of the mutational time scale. Most tumour types, including breast, ovarian and colorectal cancers, show relatively broad periods of chromosomal instability indicating a very variable timing of gains across samples.

**Figure 2.**
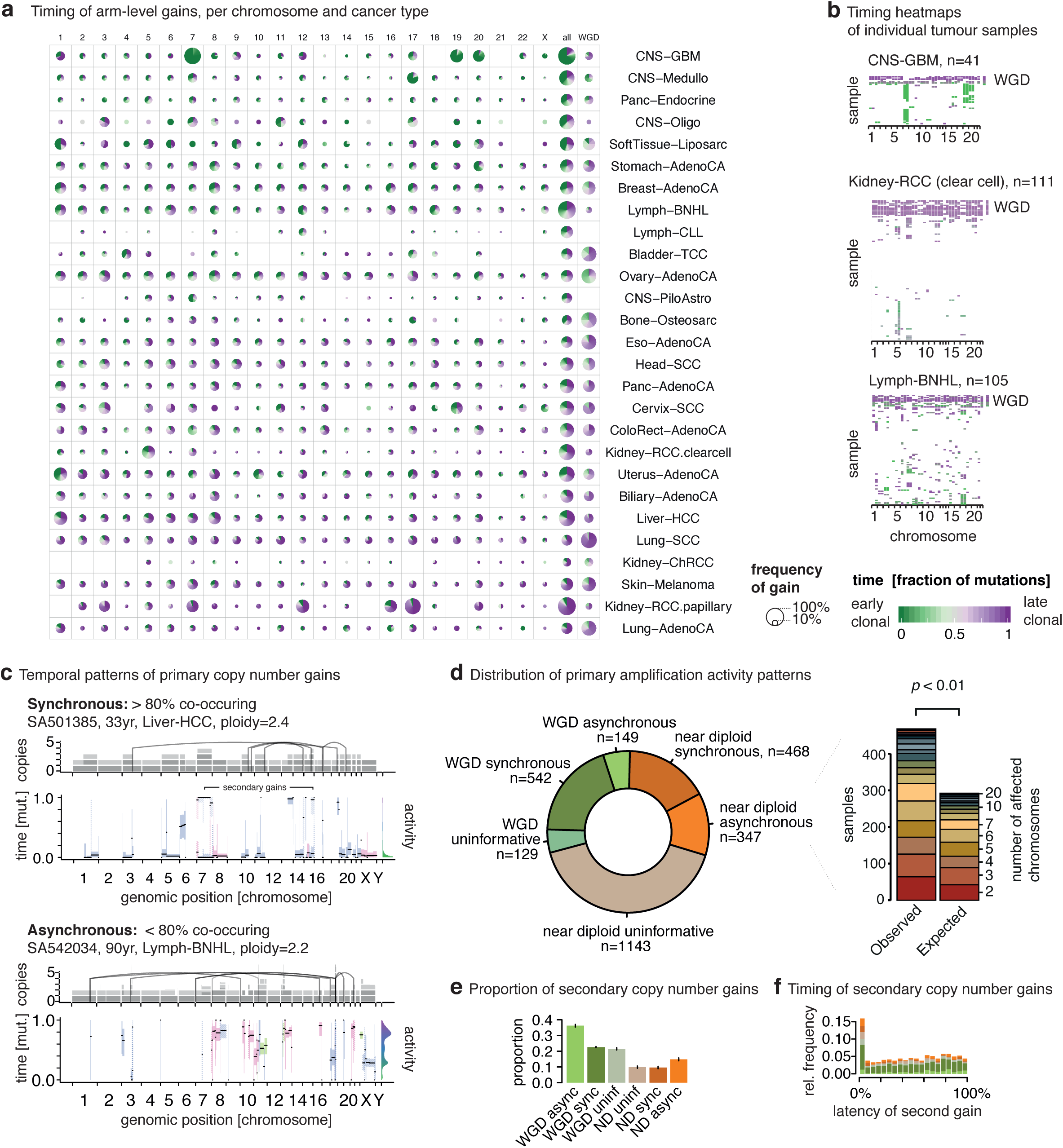
Pan-cancer timing patterns of arm-level gains. (**a**) Overview of timing arm-level copy number gains across different cancer types. Pie charts depict the distribution of the inferred mutation time for a given copy number gain in a cancer type. Green colours denote early clonal gains prior to all point mutations (mutation time t=0) with a gradient to purple for late clonal gains after all clonal point mutations (t=1). The size of each chart is proportional to the recurrence of this event in a given tissue. (**b**) Heatmaps representing timing estimates of gains on different chromosome arms (x-axis) for individual samples (y-axis) for selected tumour types. (**c**) Two near-diploid example cases illustrating synchronous gains with a single peak in amplification activity (top) and asynchronous gains with multiple amplification periods (bottom). Secondary gains leading to three copies of the same allele are separately indicated. (**d**) Distribution of synchronous and asynchronous gain patterns across samples, split by whole genome duplication status (left). Uninformative samples carry too few or too small gains to be timed accurately. Systematic permutation tests reveal a 61% enrichment of synchronous gains in near-diploid samples (right). (**e**) Proportion of large copy number segments with secondary gains per timing class. (**f**) Distribution of the relative latency of secondary gains, scaled to the time after the first gain. 0% = synchronous with first gain, 100% = at MRCA.

There are, however, certain tumour types with consistently early or late gains of specific chromosomal regions. Most pronounced is glioblastoma, where 90% of tumours harbour single copy gains of chromosome 7, 19 or 20 (**Fig. 2a-b**). Strikingly, these gains are consistently timed within the first 10% of clonal mutational time, suggesting they arise very early in a patient’s lifetime. In the case of trisomy 7, this is further supported by reports of somatic mosaicism in normal brain cells, potentially suggesting an embryonic origin^18^. Similarly, the duplications leading to isochromosome 17q in medulloblastoma are timed exceptionally early. Although less pronounced, gains of chromosome 18 in B-cell non-Hodgkin lymphoma, as well as gains of the q arm of chromosome 5 in clear cell renal cell carcinoma, often have a distinctively early timing within the first 50% of mutation time^8^. In papillary renal cell carcinoma, trisomy 7 is highly recurrent and occurs within the last 10% of clonal mutational time.

Intriguingly, we observed that gains in the same tumour often appear to occur at a similar time, pointing towards punctuated bursts of copy number gains involving the majority of gained segments (**Fig. 2c**). While this is expected in tumours with WGD (**Fig. 1b**), it may seem surprising to observe synchronous gains (defined as more than 80% of gained DNA in a single event) in near-diploid tumours. Still, synchronous gains are frequent, occurring in a striking 57% (468/815) of informative near-diploid tumours, 61% more frequently than expected by chance (*p* < 0.01, permutation test; **Fig. 2d**). Only 6% of co-amplified chromosomal segments were linked by a direct inter-chromosomal structural variant, indicating that the gains arise without being necessarily physically linked. Only 10-20% of arm-level gains increment the allele-specific copy number by more than one (**Fig. 2e**), suggesting that most of these gains arise through mis-segregation of single copies during anaphase. This notion is further supported by the observation that in about 85% of segments with two gained alleles, the second gain appears with noticeable latency after the first (**Fig. 2f**). Combined, a pattern emerges in which arm-level gains tend to arise in distinct mitotic crises that lead to numerical and sometimes structural changes of multiple chromosomes, adding (and possibly losing) one allelic copy at a time, in line with similar observations of punctuated evolution in breast cancer^19^.

### Timing of mutations in driver genes

As outlined above, point mutations (SNVs and indels) can be qualitatively assigned to different epochs, allowing the timing of driver mutations (**Fig. 1a, 3a**). Using a panel of 453 cancer driver genes^20^, we find that pathogenic mutations in the 50 most common drivers are predominantly clonal, and often early clonal (**Fig. 3a-b**). For example, driver mutations in *TP53* and *KRAS* are 12x and 8x enriched in early clonal stages, respectively. For *TP53*, this trend is independent of tumour type (**Fig. 3c**). Mutations in *PIK3CA* are 2x more frequently clonal than subclonal, while non-coding changes near the *TERT* gene are 3x more frequently early clonal than expected.

**Figure 3.**
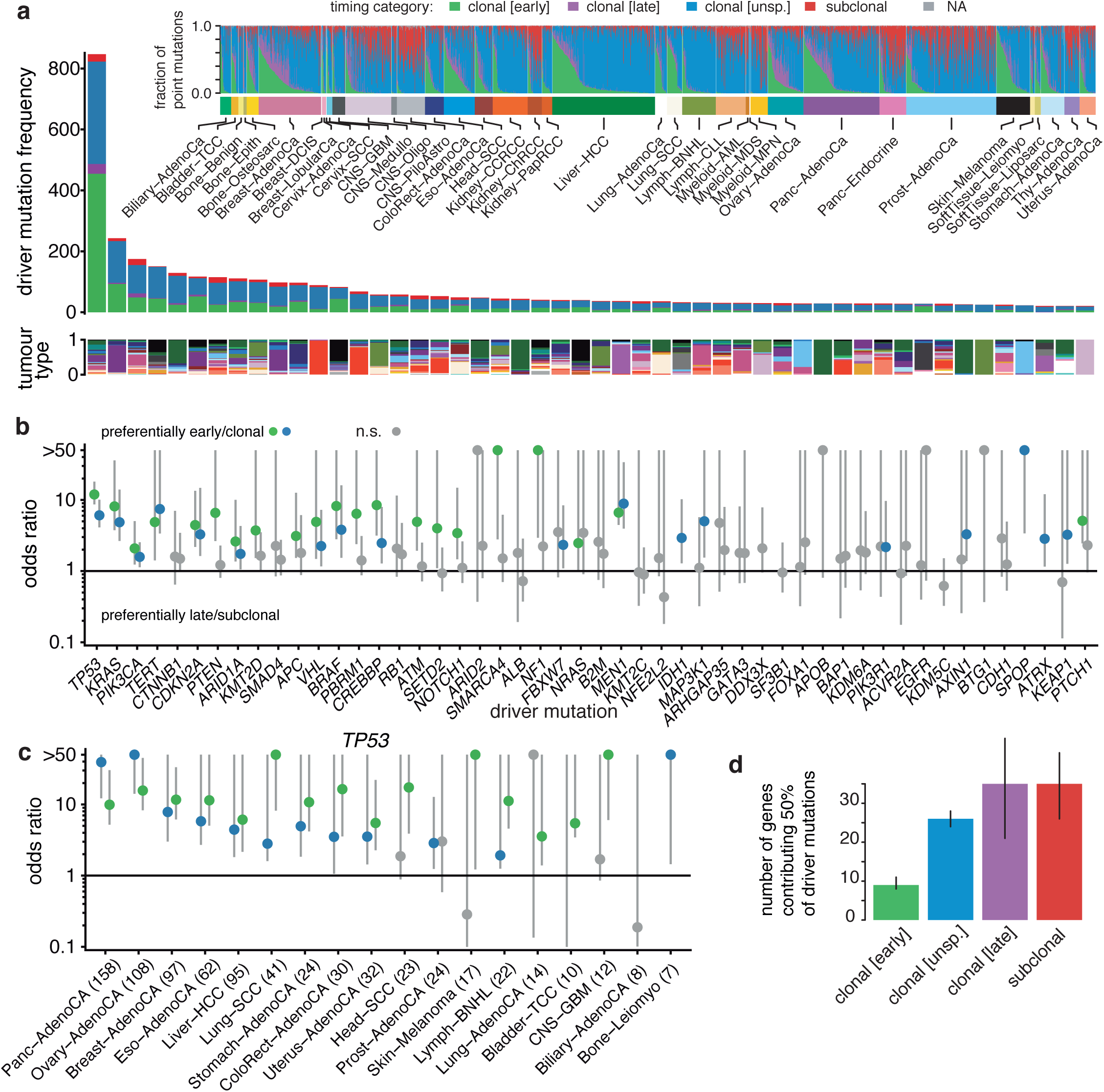
Timing of driver point mutations. (**a**) Top: distribution of point mutations over different mutation periods in 2,778 samples. Middle: timing distribution of driver mutations in the 50 most recurrent lesions across 2,583 white listed samples from unique donors. Bottom: distribution of driver mutations across cancer types; colour as defined in the inset. (**b**) Relative timing of the 50 most recurrent driver lesions, calculated as the odds ratio of early versus late clonal driver mutations versus background (green, purple) or clonal versus subclonal. Odds ratios overlapping 1 in less than 5% of bootstrap samples are considered significant and have been coloured. Odds ratios have been truncated at 50. (**c**) Relative timing of *TP53* mutations across cancer types, coloured and truncated as in (**b**). The number of mutations in parenthesis represents the total number of *TP53* mutations used in the analysis for each cancer type. (**d**) Estimated number of unique lesions (genes) contributing 50% of all driver mutations in different timing epochs. Error bars denote the range between 0 and 1 pseudocounts.

Overall, common driver mutations predominantly occur during the early clonal period of tumour evolution. To understand how the entire landscape of all 453 driver genes changes over time, we calculated how the number of driver mutations relates to the number of driver genes in each of the evolutionary stages. This reveals an increasing diversity of driver genes mutated at later stages of tumour development: 50% of all early clonal driver mutations occur in only 9 genes, whereas the corresponding proportion of late and subclonal mutations occur in approximately 35 different genes each, a nearly 4-fold increase (**Fig. 3d**). These results are consistent with previous findings in non-small-cell lung cancers^21^, and suggest that, across cancer types, the very early carcinogenic events occur in a constrained set of common drivers, while a more diverse array of drivers is involved in late tumour development.

### Relative timing of somatic driver events

Next, we sought to better understand the sequence and timing of events during tumour evolution by integrating the timing of driver point mutations and recurrent copy number changes across cancer samples. We calculated an overall probabilistic ranking of lesions (**Fig. 4a**), detailing whether each mutation occurs preferentially early or late during tumour evolution, by aggregating single patient partial order trajectories (**Fig. 4b-d**) within each cancer type (**Supplementary Information**, section 3.2, **Extended Data Fig. 2-4**).

**Figure 4.**
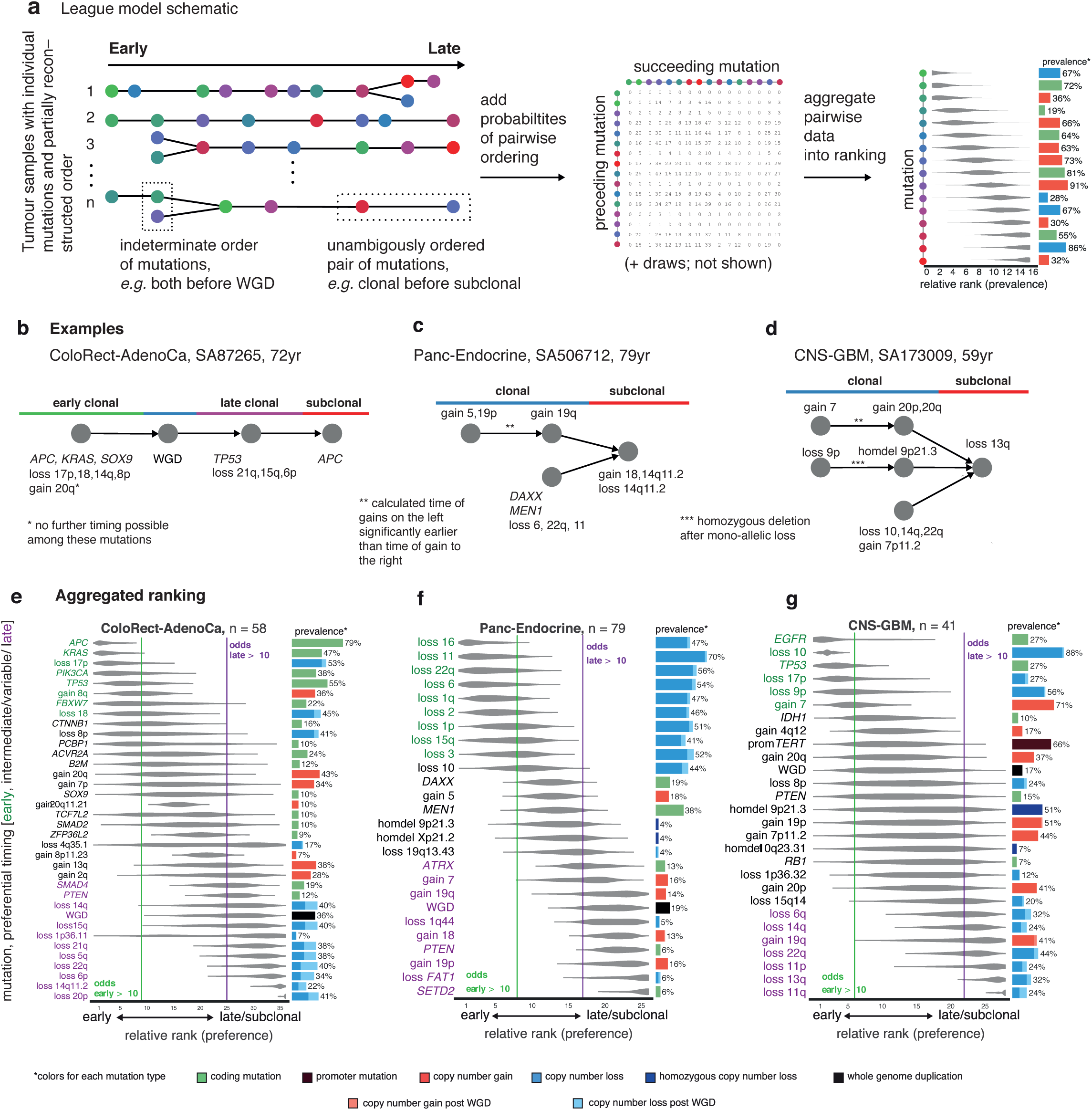
Relative ordering of somatic events. Preferential ordering of somatic copy number events and driver point mutations ithin tumour cohorts. (**a**) Schematic representation of the ordering process. (**b-d**) examples of individual patient trajectories (partial ordering relationships), the constituent data for the League model ordering process. Preferential ordering diagrams for (**e**) colorectal adenocarcinoma, (**f**) pancreatic neuroendocrine cancer and (**g**) glioblastoma. Probability distributions show the uncertainty of timing for specific events in the cohort. Events with odds above 10 (either earlier or later) are highlighted. The prevalence of the event type in the cohort is displayed as a bar plot on the right.

We have confirmed through simulations that the average order of events is correctly estimated through aggregating individual patient trajectories (**Extended Data Fig. 3**), which can be of varying level of detail depending on the sample (**Fig. 4a**, **Supplementary Information,** section 3.2). As seen from example patient tumours (**Fig. 4b-d**), whole genome sequencing provides insights on the actual order of driver mutations, even on the single-sample level (*e.g., APC* and *KRAS* mutations occur before events in *TP53* in colorectal tumour SA87265; **Fig. 4b**). Through aggregation of many such partial order relationships, we can infer probabilistic rankings of events within each cancer type (**Fig. 4e-g**, **Extended Data Fig. 1**).

In colorectal adenocarcinoma, for example, we find *APC* mutations to have the highest odds of occurring early, followed by *KRAS,* loss of 17p and *TP53,* and *SMAD4* (**Fig. 4e**). Whole-genome duplications have an intermediate ranking, indicating a variable timing, while many chromosomal gains and losses are typically late. These results are in agreement with the classical progression *APC-KRAS-TP53* model of Vogelstein and Fearon^22^, but add considerable detail.

In other cancer types, the sequence of events in cancer progression has not previously been studied in as much detail. For example, in pancreatic neuroendocrine cancers, we find that many chromosomal losses, including those of chromosomes 2, 6, 11 and 16, occur early, followed by driver mutations in *MEN1* and *DAXX* (**Fig. 4f**). WGD events occur late, after many of these tumours have reached a pseudo-haploid state due to wide-spread chromosomal losses. In glioblastoma, we find that loss of chromosome 10 and driver mutations in *TP53* and *EGFR* are very early, often preceding early gains of chromosomes 7, 19 and 20 (as described above) (**Fig. 4g**). *TERT* promoter mutations tend to occur at early to intermediate time points, while other driver mutations and copy number changes tend to be later events.

Across cancer types, we typically find *TP53* mutations early, as well as losses of chromosome 17 (**Extended Data Fig. 1**). WGD events usually have an intermediate ranking and the majority of copy number changes occur after WGD. We also find that losses typically precede gains, and consistent with the results above, common drivers typically occur before rare drivers.

### Timing of mutational signatures

The cancer genome is shaped by various mutational processes over its lifetime, stemming from exogenous and cell-intrinsic DNA damage and error-prone DNA replication, each leaving behind characteristic mutational spectra, termed mutational signatures^23,24^. To quantify the changing activity of mutational processes operating in each cancer genome, we assessed differences of the observed mutational spectra between early clonal, late clonal and subclonal epochs (**Supplementary Information**). As shown in **Fig. 5a**, there are indeed examples of striking changes between early and late clonal single base substitution spectra, indicating that the underlying mutational processes have considerably changed during the clonal life history of these cancers. Overall, we find evidence for a changing mutational spectrum between early and late clonal time points in 29% (530/1,852) of informative samples (*p* < 0.05, Bonferroni-adjusted likelihood-ratio test), typically changing 19% of mutations (median absolute difference; range 4%–66%).

**Figure 5.**
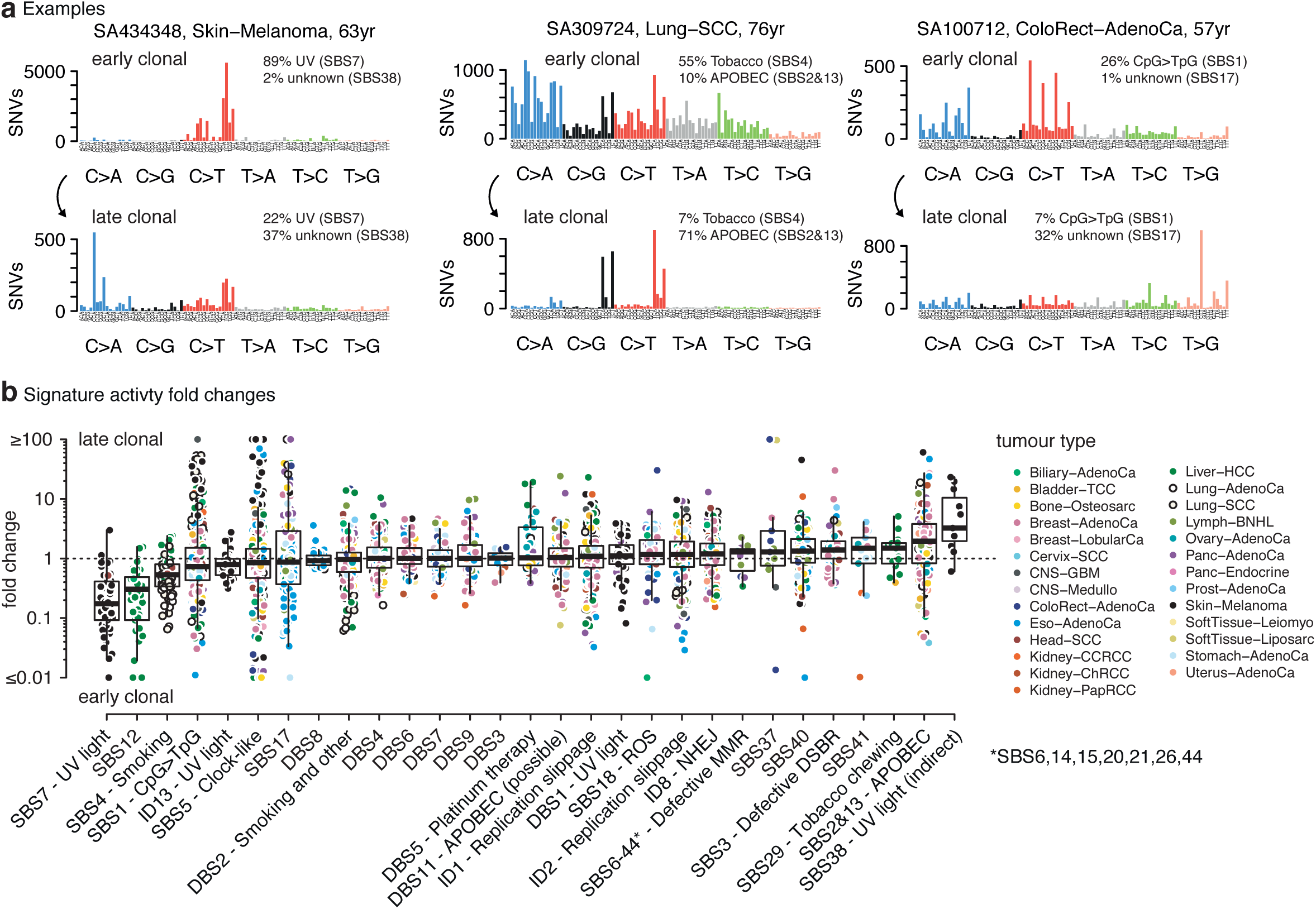
Dynamic mutational processes during early and late clonal tumour evolution. **(a)** Examples of tumours with substantial changes between mutation spectra of early and (top) late clonal time points (bottom). or each sample, the attribution of mutations to the most characteristic single-base signatures are shown. Percentages missing from 100% are due to other, less characteristic contributions. **(b)** Fold changes between relative proportions of early and late clonal mutations attributed to individual mutational signatures. Points are coloured by tissue type. Data are shown for n = 530 samples with measurable changes in their overall mutation spectra and restricted to signatures active in at least 10 samples.

To quantify whether the observed temporal changes can be attributed to known mutational processes, we decomposed the mutational spectra at each timepoint into a catalogue of 57 mutational signatures, tailored to this cohort and fully described in Alexandrov *et al.*^24^ **(Fig. 5a; Methods)**. To this end, we were also able to make use of two new categories of signatures: a series of doublet base substitution signatures, and a series of indel signatures^24^.

The fold-changes of mutations attributed to a given signature between early and late clonal stages of tumour evolution indeed vary by several orders of magnitude, revealing dynamic evolution of many mutational processes during the life history of many, but not all tumours (**Fig. 5b**).

In addition to this predominantly undirected temporal variability, a number of signatures demonstrate directional trends in their activity over time (**Fig. 5c**). As one may expect, we observe signatures of exogeneous mutagens to be predominant in the early clonal stages of tumourigenesis. These include tobacco smoking (single base signature SBS4), median fold change 0.54, IQR [0.34, 0.8]), which is most pronounced in lung adenocarcinoma (median change 0.32; IQR [0.2,0.37]; **Extended Data Fig. 5**) and consistent with previous reports^25,26^. Quite similarly, the signature of UV light in melanoma contributes far fewer mutations at late clonal stages (SBS7; mean fold change 0.17, IQR [0.09, 0.41]). Another strong decrease over time, similar to that of tobacco smoke and UV damage, is found for a signature of unknown aetiology, SBS12, which acts mostly in liver cancers (median fold change 0.31, IQR [0.09, 0.49]).

Some mutational processes are preferentially active in late stages of clonal evolution. For example, we see that APOBEC mutagenesis (SBS2 and SBS13) increases in many cancer types from the early to late clonal stages (median fold change 2, IQR [0.83, 3.8]), as does a newly described signature attributed to indirect damage from UV light (SBS38; median fold 3.2, IQR [2.0,10])^24^. Lastly it is interesting to note that chemo-resistant ovarian cancers display an increase in the mutational signature associated with platinum-based therapy (double base signature DBS5; median fold change 1.8, IQR [1.0, 3.3]).

### Chronological time estimates of whole genome duplications and subclonal diversification

Changes in the mutation rate of cancers influence timing estimates based upon mutational data. Due to increased proliferation and in some cases acquired hypermutation, one would generally expect an increase in the mutation rate over time, yet some mutational processes appear more variable than others.

CpG>TpG mutations as the result of spontaneous deamination of methylated cytosines (SBS1), constitute a promising candidate for a clock-like mutational process^27^, as they are ubiquitously found in all cancer and normal tissues^28,29^. The number of CpG>TpG mutations increase constantly with age in this cohort (**Fig. 6a**), with moderate variation between samples and across tissues (**Extended Data Fig. 6a**). As long as the rate of the mutational clock is constant within a sample, any inter-sample variation can be accounted for, since the age at diagnosis is used to calibrate the mutation clock in each tumour.

**Figure 6.**
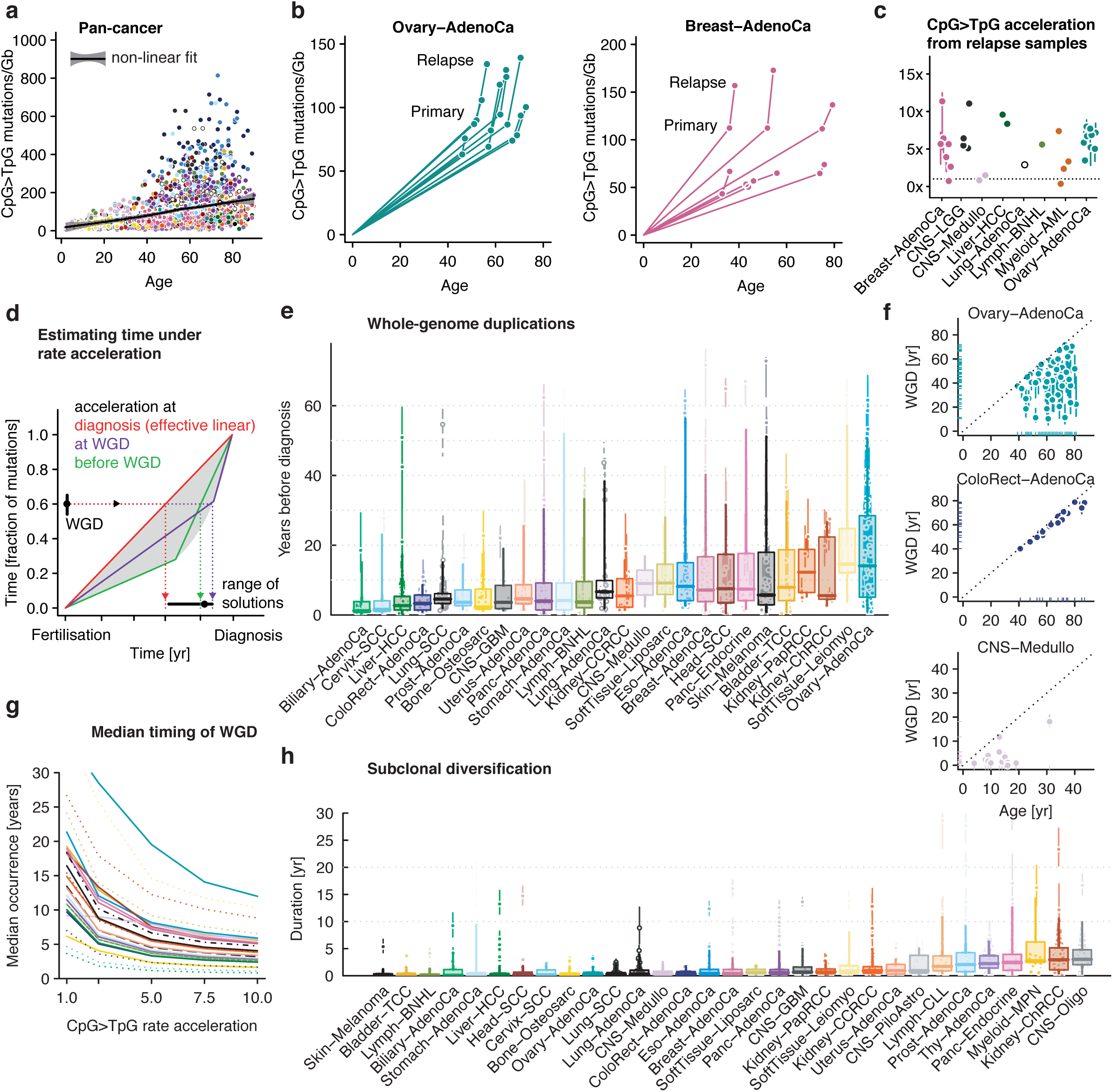
Real-time estimation of mutational landmarks. (**a**) CpG>TpG mutations/Gb (scaled to the effective genome size) as a function of age at diagnosis across cancer types. Colours are defined in panel **h**. **(b**) CpG>TpG mutations/Gb for ovarian and breast cancer samples with matched primary and relapse samples. (**c**) CpG>TpG mutation rate increase inferred from paired samples for 6 cancer types. (**d**) Schematic of absolute timing under mutation rate increase. (**e**) Time of occurrence of WGD in individual patients, split by tumour type. The error bars at individual data points are 80% confidence intervals, reflecting uncertainty stemming from the number of mutations per segment and onset of the rate increase, which was chosen between 2.5x and 7.5x. Overlaid boxplots demarcate the quartiles and median of the distribution with whiskers indicating 5% and 95% quantiles. (**f**) Scatter plots showing the time of diagnosis (x-axis) and inferred time of WGD (y-axis) with error bars as in **e**. (**g**) Median timing of WGD per cancer type, as a function of CpG>TpG rate increase. Dotted lines indicate cancer types with less than 5 informative samples. Dashed and dash-dotted lines represent Lung-AdenoCa and Lung-SCC, respectively; colours as in part **e.** (**h**) Timing of subclonal diversification using CpG>TpG mutations in individual patients.

However, there may also be an increase in the CpG>TpG rate over time, which can be estimated in two ways. First, sequencing data of matched primary and relapse samples from the same donor suggest that there is an increased rate of mutation accumulation between diagnosis and relapse (as evidenced here for publicly available data from ovarian and breast cancer^30,31^), **Fig. 6b**, and therefore that this acceleration likely extends to periods immediately prior to diagnosis. Formalising this argument for eight cancer types^30-33^ yields a typical acceleration of around 5x [range: 1x-12x] post-diagnosis relative to pre-diagnosis (**Fig. 6c**). A second line of evidence comes from a systematic analysis of the relation between CpG>TpG mutation burden and age at diagnosis^34^ in this cohort (**Extended Data Fig. 6a)**. This indicates that, across cancer types, typically 20-40% of mutations are found independent of age and are thereby likely caused by an acceleration during cancer progression (**Extended Data Fig. 6b-c**). This constant offset is broadly compatible with a 5x acceleration over a period of about five years prior to a diagnosis at a typical age of 60 years.

One effect of an increasing mutation rate is that chronological time estimates appear closer to diagnosis and have higher variance, as the exact onset of acceleration is unknown (**Fig. 6d**). Applying this logic to the timing of whole-genome duplications based on CpG>TpG mutations, we find that the typical timing of WGD is about one decade before diagnosis (**Fig. 6e**). However, there is substantial variability among samples of a given tumour type, with many cases dating back more than two decades. This latency corresponds to rather different patterns in the lifetime of individuals (**Fig. 6f**): Ovarian adenocarcinomas have WGD occurring more than two decades before diagnosis in almost half of patients. For these early cases, the estimated age of WGD falls well into female reproductive life, concordant with the observation that the number of ovulations is a risk factor for ovarian cancer^35^. Conversely, for colorectal adenocarcinoma, most duplications shortly precede diagnosis. Finally, in the childhood cancer medulloblastoma, the majority of WGD events fall into the first two years of life, suggesting an early developmental onset of the disease.

Reassuringly, the earliness of WGDs is caused by depletion of early (that is co-amplified) mutations and is independent of the overall mutation burden (**Extended Data Fig. 6d-f**), making it unlikely that the observed early duplications are artefacts of acquired hypermutation. While the observation that the majority of CpG>TpG mutations occur proportionally to age places some bounds on the strength of acceleration, some uncertainty persists. We thus note that, without any acceleration, the estimated median latency between WGD and diagnosis would be 6-35 years for the majority of cancer types analysed in this study; conversely at an acceleration of 10x the median latency would be 1.6-12 years (range for tumour types with ≥10 samples; **Fig. 6g**).

We used a similar approach to calculate the duration of subclonal diversification, *i.e.* the time between the MRCA and the last detectable subclone. The typical timing is considerably shorter and about 1yr, but interestingly, there are also cases where this period is estimated to last more than ten years (**Fig. 6h**). We note, however, that timing of the subclonal diversification period is more difficult and the results presented here are lower bounds. This is because there may be earlier subclonal branches missed in a single sample, there is limited power to detect very late and deeply subclonal branches and also because the underlying phylogenetic relationships between subclones cannot always be unambiguously resolved and were treated as densely branching. Assuming linear instead of branching phylogenies approximately doubles the inferred latency (**Extended Data Fig. 6g**).

While the exact timing of individual samples remains challenging due to low CpG>TpG mutation numbers and unknown mutation rates for individual tumours, these findings are entirely consistent with epidemiological observations: cancer generally arises past the age of 50 (Ref. ^36^), and the typical latency between carcinogen exposure and cancer detection, e.g. in tobacco-associated cancers, is several years to multiple decades^37^. Furthermore, the progression of most known precancerous lesions to carcinomas usually spans multiple years, if not decades^38-45^. Our data corroborate these time scales and show that they hold in cases without detectable premalignant conditions, raising hopes that these tumours could also be detected in precancerous stages.

## Discussion

Taken together, the data presented here enable us to draw semi-quantitative timelines summarising the typical evolutionary history of each cancer type (**Fig. 7**). These make use of the qualitative timing of point mutations and copy number drivers, as well as signature activities, which can be interleaved with the chronological estimates of WGD and MRCA (**Fig. 7**, **Extended Data Fig. 1** for a complete list of all cancer types).

**Figure 7.**
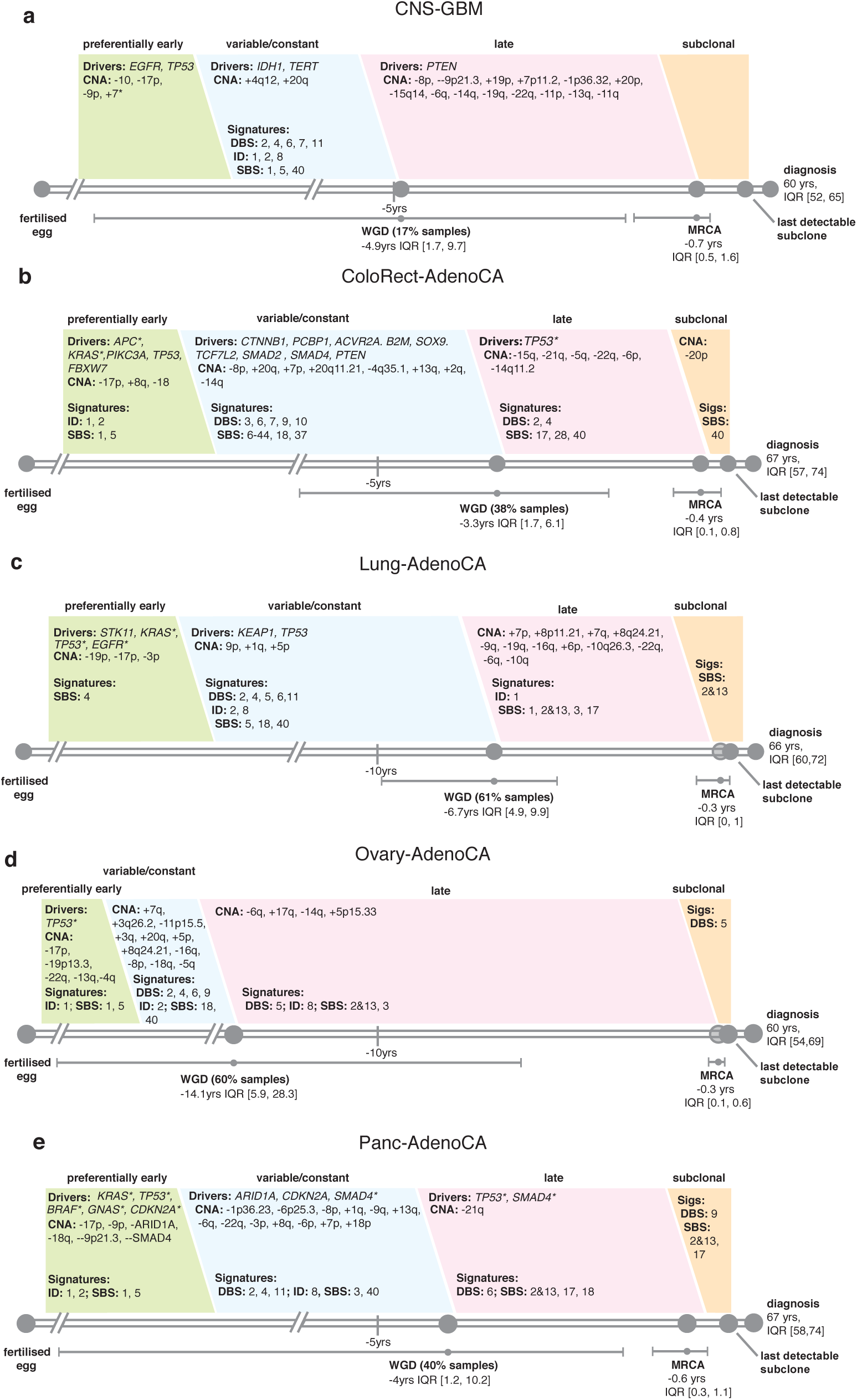
Cancer timelines. Typical timelines of tumour development, for (**a**) glioblastoma, (**b**) colorectal adenocarcinoma, (**c**) squamous cell lung cancer, (**d**) ovarian adenocarcinoma, and (**e**) pancreatic adenocarcinoma. Each timeline represents the length of time, in years, between the fertilised egg and the median age of diagnosis per cancer type. Real time estimates for major events, such as WGD and the emergence of the MRCA are used to define early, variable, late and subclonal stages of tumour evolution approximately in chronological time. Driver mutations and copy number aberrations are shown in each stage according to their preferential timing, as defined by relative ordering. Mutational signatures that, on average, change over the course of tumour evolution (confidence intervals not overlapping 0) are shown in the epoch in which their activity is greatest. Where applicable, lesions with a timing already defined in the literature are annotated, * denotes events that agree with our timing, while † denotes events that either were not found in our study, or were found to have a different timing.

Individual tumour types show characteristic evolutionary trajectories, reflecting differences in the mutational processes driving evolution, the portfolio of driver mutations and their timing. While glioblastoma, for example, is often initiated by mutations in *EGFR* and gain of chromosome 7, *APC* and *KRAS* mutations are typically the earliest events in colorectal adenocarcinoma, followed by *TP53*, in line with classic findings^22^. This is similar to reports of frequent *KRAS* and *TP53* mutations and deletions of 17p in precursors to pancreatic adenocarcinoma^46^, confirming the early role found here. *TP53* with accompanying 17p deletion is one of the most frequent initiating mutations in a variety of cancers, including ovarian cancer, where it is the hallmark of its precancerous precursor lesion^47^. CpG>TpG mutations (SBS1) are active throughout tumour development, however, their relative contribution decreases in favour of more diverse mutational processes acting at later stages of cancer development. Similarly, genomic instability, leading to frequent gains and losses, is typically seen late in cancer progression. Unlike most other cancers, high grade serous ovarian adenocarcinomas typically acquire chromosomal gains within the first half of clonal evolution (**Fig. 7d**). Our findings are consistent with these tumours having a high frequency of *TP53* and homologous recombination repair defects^48^, and rendering them among the most genomically unstable of all solid cancers^49^.

Across cancer types, our findings provide insight into how selection acts during cancer development, with an increased repertoire of driver mutations in late tumour evolution. Genetic canalization in early stages suggests that epistasis of fitness effects is constraining evolution to select for only a small set of mutations in genes and chromosomal aberrations that are able to initiate neoplastic transformation. Over time, as tumours evolve, the small- and large-scale somatic changes they subsequently accumulate propel them towards increasingly diverse paths driven by individually rare, atypical driver mutations.

Our analysis reveals that tumour evolution is driven by crises leading to simultaneous gains of multiple or even all chromosomal segments, likely as the result of numerical and structural misconfiguration during a single mitotic failure. Therefore, the extent of chromosome-scale copy number aberration observed in tumour genomes is likely to be caused by much fewer events than the number of chromosomes affected.

Finally, our study enlightens the typical time scales of *in vivo* tumour progression, with WGD occurring typically a decade before diagnosis and the onset of subclonal variegation a few years before diagnosis. The presence of genetic aberrations for 10 years raises hopes that aberrant clones could be detected early, prior to reaching fully malignant potential. Interestingly, this also shows that individual mutations, including driver gene mutations and chromosomal gains, can occur several decades before cancer diagnosis, demonstrating how cancer genomes have been shaped by a lifelong process of somatic evolution.

## Methods

### Data set

The PCAWG series consists of 2,778 tumour samples (2,703 white listed, 75 grey listed) that represent 2,658 cancers. All samples in this dataset underwent whole genome sequencing (minimum average coverage 30x in the tumour, 25x in the normal), and were processed with a set of project-specific pipelines for alignment, variant calling, and quality control^15^. Copy number calls were established by combining the output of six individual callers into a consensus using a multi-tier approach, resulting in a copy number profile, a purity and ploidy value and whether the tumour has undergone a whole genome duplication^50^. Consensus subclonal architectures have been obtained by integrating the output of eleven subclonal reconstruction callers^50^, after which all SNVs, indels and SVs are assigned to a mutation cluster using the MutationTimer.R approach (**Supplementary Information**). Driver calls have been defined by the PCAWG Driver working group^20^ while mutational signatures are defined by the PCAWG Signatures working group^24^. A more detailed description can be found in **Supplementary Information**, section 0.

### Timing of gains

We used three related approaches to calculate the timing of copy number gains (see **Supplementary Information**, section 1). In brief, the common feature is that the expected variant allele frequency of a mutation is related to the underlying number of alleles carrying a mutation according to the formula

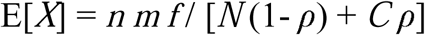

Here *X* is the number of reads, *n* denotes the coverage of the locus, the mutation copy number *m* is the number of alleles carrying the mutation (which is usually inferred), *f* is the frequency of the clone carrying the given mutation (*f* = 1 for clonal mutations). *N* is the normal copy number (2 on autosomes, 1 or 2 for chromosome X and 0 or 1 for chromosome Y), *C* the total copy number of the tumour and *ρ* the purity of the sample.

The number of mutations at each allelic copy number then informs about the time when the gain has occurred. The basic formulae for timing each gain are, depending on the copy number configuration:

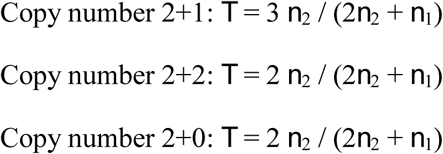

Here 2+1 refers to major and minor copy number of 2 and 1, respectively. Methods differ slightly in how the number of mutations present on each allele are calculated and how uncertainty is handled (Supplementary Informatio006E).

### Timing of mutations

The mutation copy number *m* and the clonal frequency *f* is calculated according to the principles indicated above. Details can be found in **Supplementary Information**, section 1.2. Mutations with *f* = 1 are denoted as *clonal*, and mutations with *f* < 1 as *subclonal*. Mutations with *f* = 1 and *m* > 1 are denoted as *early clonal* (coamplified). In cases with *f* = 1, *m* = *1* and *C* > 2, mutations were annotated as *late clonal*, if the minor copy number was 0, otherwise *clonal* [*unspecified*] (**Supplementary Information**, section 1.2.)

### Timing of driver mutations

A catalogue of driver point mutations (SNVs and indels) was provided by the PCAWG Drivers and Functional Interpretation Group^20^. The timing category was calculated as above. From the four timing categories, odds ratios of early/late clonal and clonal (early, late or unspecified clonal)/subclonal were calculated for driver mutations against the distribution of all other mutations present in fragments with the same copy number composition in the samples with each particular driver. The background distribution of these odds ratios was assessed with 1,000 bootstraps (**Supplementary Information**, section 3.1.)

### Integrative timing

For each pair of driver point mutations and recurrent copy number alterations, an ordering was established (earlier, later or unspecified). The information underlying this decision was derived from the timing of each driver point mutation, as well as from the timing status of clonal and subclonal copy number segments. These tables were aggregated across all samples and a sports statistics model was employed to calculate the overall ranking of driver mutations. A full description is given in **Supplementary Information**, section 3.2.

### Timing of mutational signatures

Mutational trinucleotide substitution signatures, as defined by the PCAWG Mutational Signatures Working Group^24^, were fit to samples with observed signature activity, after splitting point mutations into either of the four epochs. A likelihood ratio test based on the multinomial distribution was used to test for differences in the mutation spectra between time points. Time-resolved exposures were calculated using non-negative linear least squares. Full details are given in **Supplementary Information**, section 4.

### Real-time estimation of WGD and MRCA

CpG>TpG mutations were counted in an NpCpG context, except for Skin-Melanoma, where CpCpG and TpCpG were excluded due to the overlapping UV mutation spectrum. For visual comparison, the number of mutations was scaled to the effective genome size, defined as the 1 / mean (*m*_*i*_ */ C*_*i*_) where *m*_*i*_ is the estimated number of allelic copies of each mutation and *C*_*i*_ the total copy number at that locus, thereby scaling to the final copy number and the time of change.

A hierarchical Bayesian linear regression was fit to relate the age at diagnosis to the scaled number of mutations, ensuring positive slope and intercept through a shared gamma distribution across cancer types.

For tumours with multiple time points, the set of mutations shared between diagnosis and relapse (*n*_D_) and those specific to the relapse (*n*_R_) was calculated. The rate acceleration was calculated as *a* = *n*_R_ / *n*_D_ × *t*_D_ / *t*_R_. This analysis was performed separately for all substitutions and for CpG>TpG mutations.

Based on these analyses, a typical increase of 5x for most cancer types was chosen, with a lower value of 2.5x for brain cancers and a value of 7.5x for ovarian cancer.

The correction for transforming an estimate of a copy number gain in mutation time into chronological time depends not only on the rate acceleration, but also on the time at which this acceleration occurred. As this is generally unknown, we performed Monte Carlo simulations of rate accelerations spanning an interval of 15 years prior to diagnosis, corresponding roughly to 25% of time for a diagnosis at 60 years of age, noting that a 5x rate increase over this duration yields an offset of about 33% of mutations, compatible with our data. Subclonal mutations were assumed to occur at full acceleration. The proportion of subclonal mutations was divided by the number of identified subclones, thus conservatively assuming branching evolution. Full details are given in **Supplementary Information**, section 5.

### Cancer timelines

The results from each of the different timing analyses are combined in timelines of cancer evolution for each tumour type (**Figure 7** and **Extended Data Fig. 1**). Each timeline begins at the fertilised egg, and spans up to the median age of diagnosis within each cohort. Real time estimates for WGD and the MRCA act as anchor points, allowing us to roughly map the four broadly defined time periods (early clonal, intermediate, late clonal and subclonal) to chronological time during a patient’s lifespan. Specific driver mutations or copy number alterations can be placed within each of these time frames based on their ordering from the league model analysis. The most active mutational signatures within each cohort are displayed, with font sizes indicating the mean proportion of mutations attributed to each signature during a given time period. Full details are given in **Supplementary Information**, section 6.

## Data availability

Raw variant calls will be made available through the PCAWG data portal, https://dcc.icgc.org/pcawg. Processed data from the PCAWG Evolution and Heterogeneity working group, including timing estimates for all copy number gains, real time estimates of WGD and MRCA, as well as mutation signature activities, is available at www.synapse.org/#!Synapse:syn14193595. Analysis code can be downloaded via the github repository https://github.com/PCAWG-11/Evolution, containing submodules reflecting software tools and analysis workflows used in this analysis.

## Extended Data Figure Legends

**Extended Data Figure 1.**
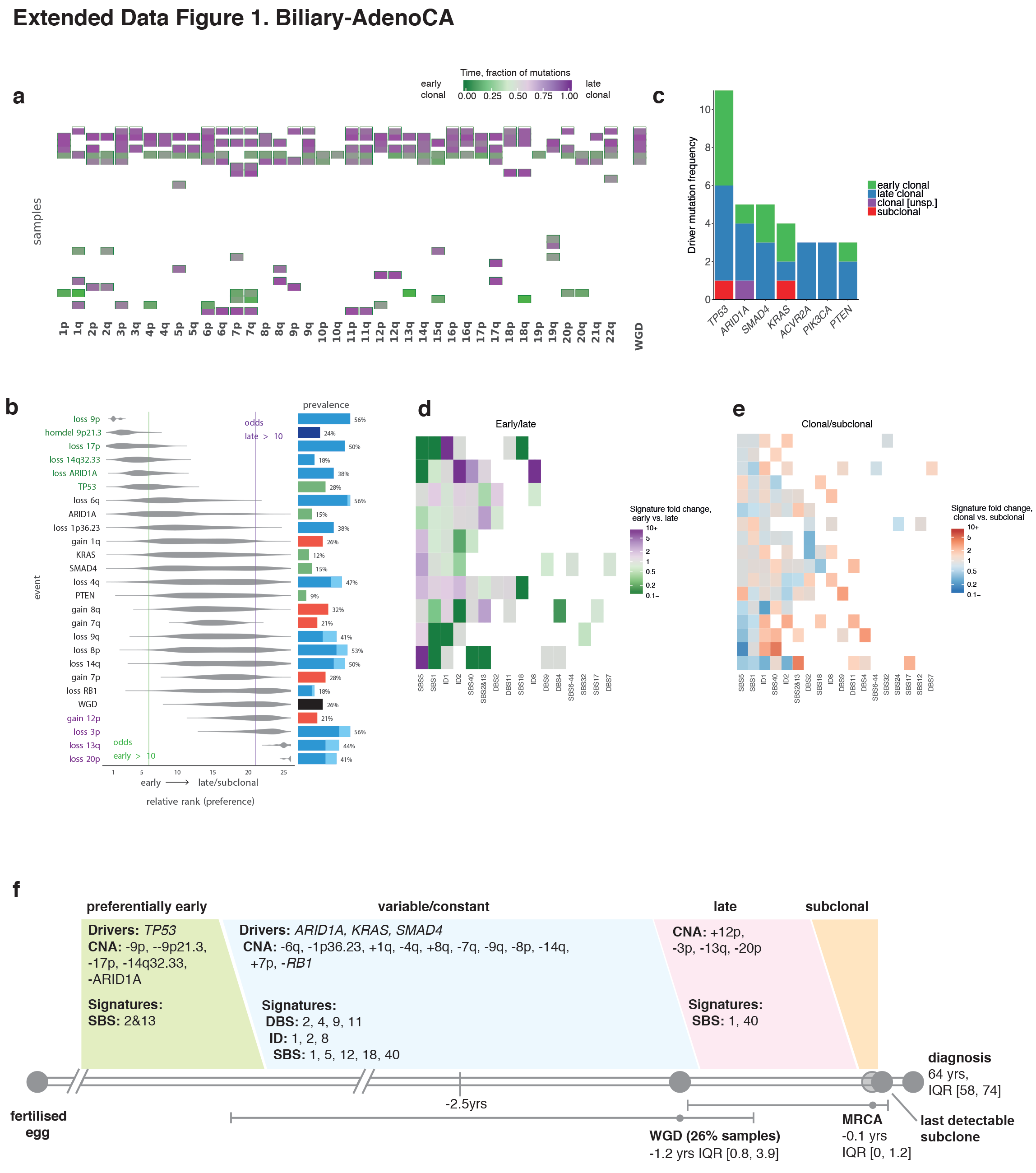
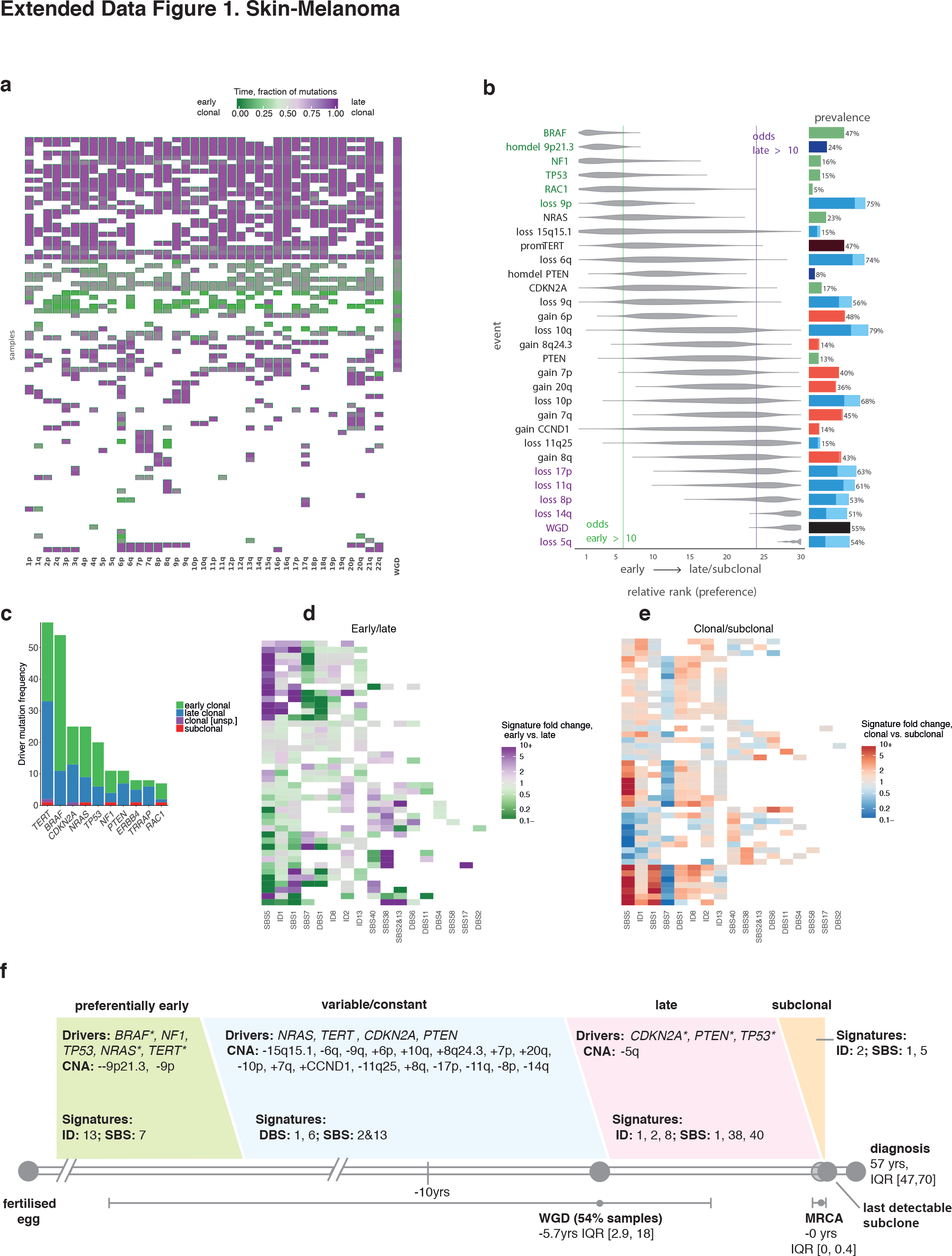
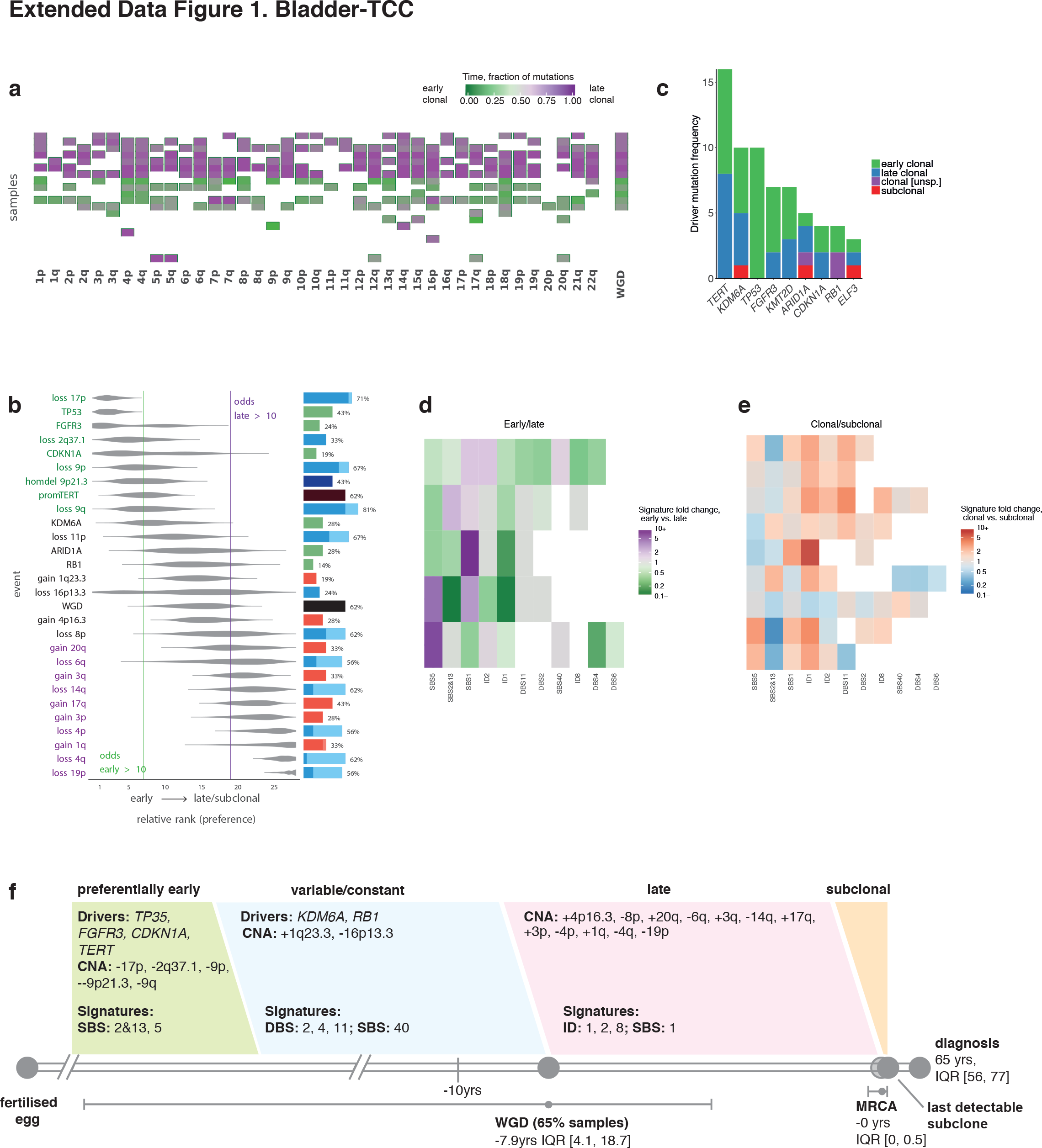
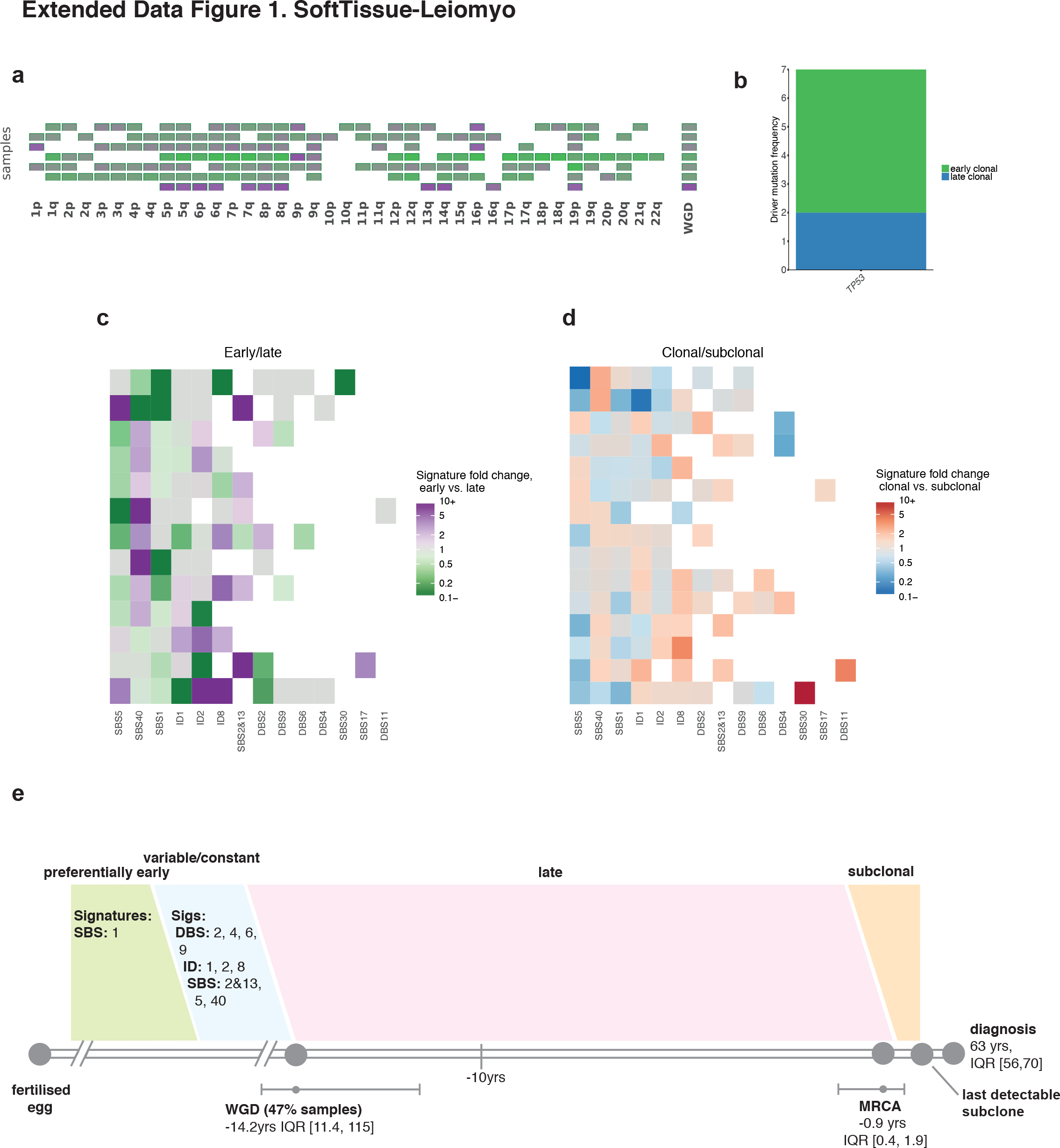
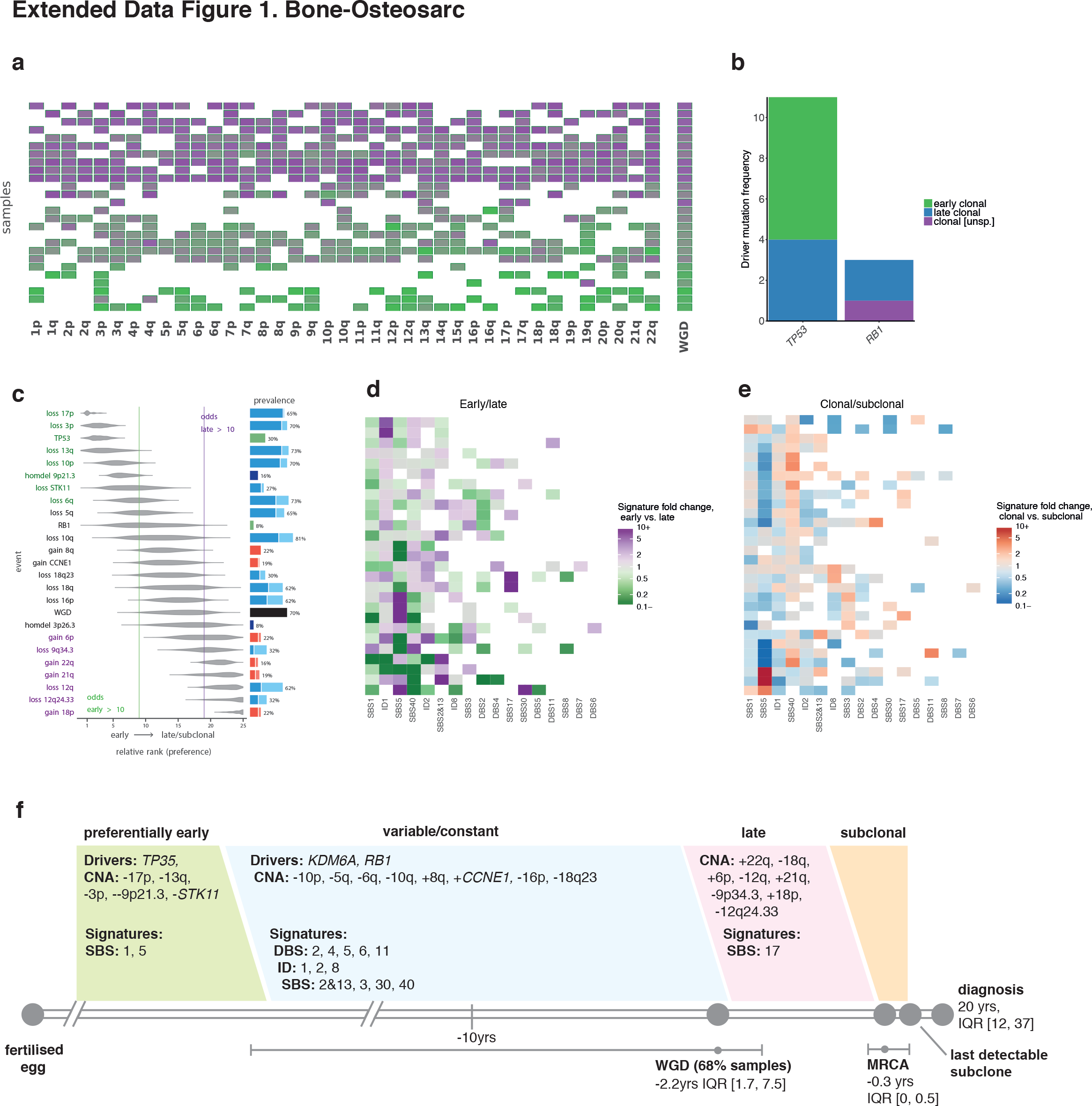
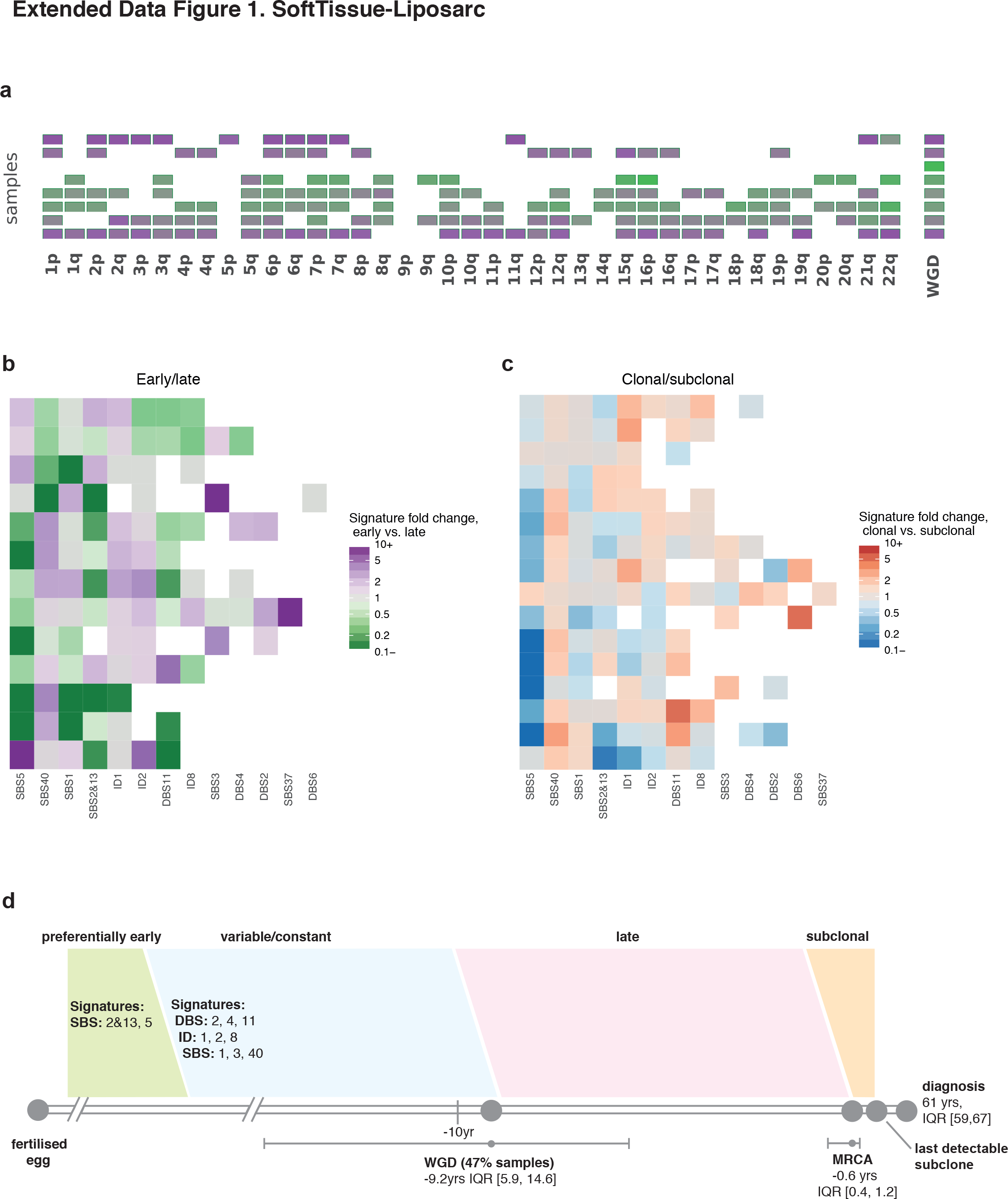
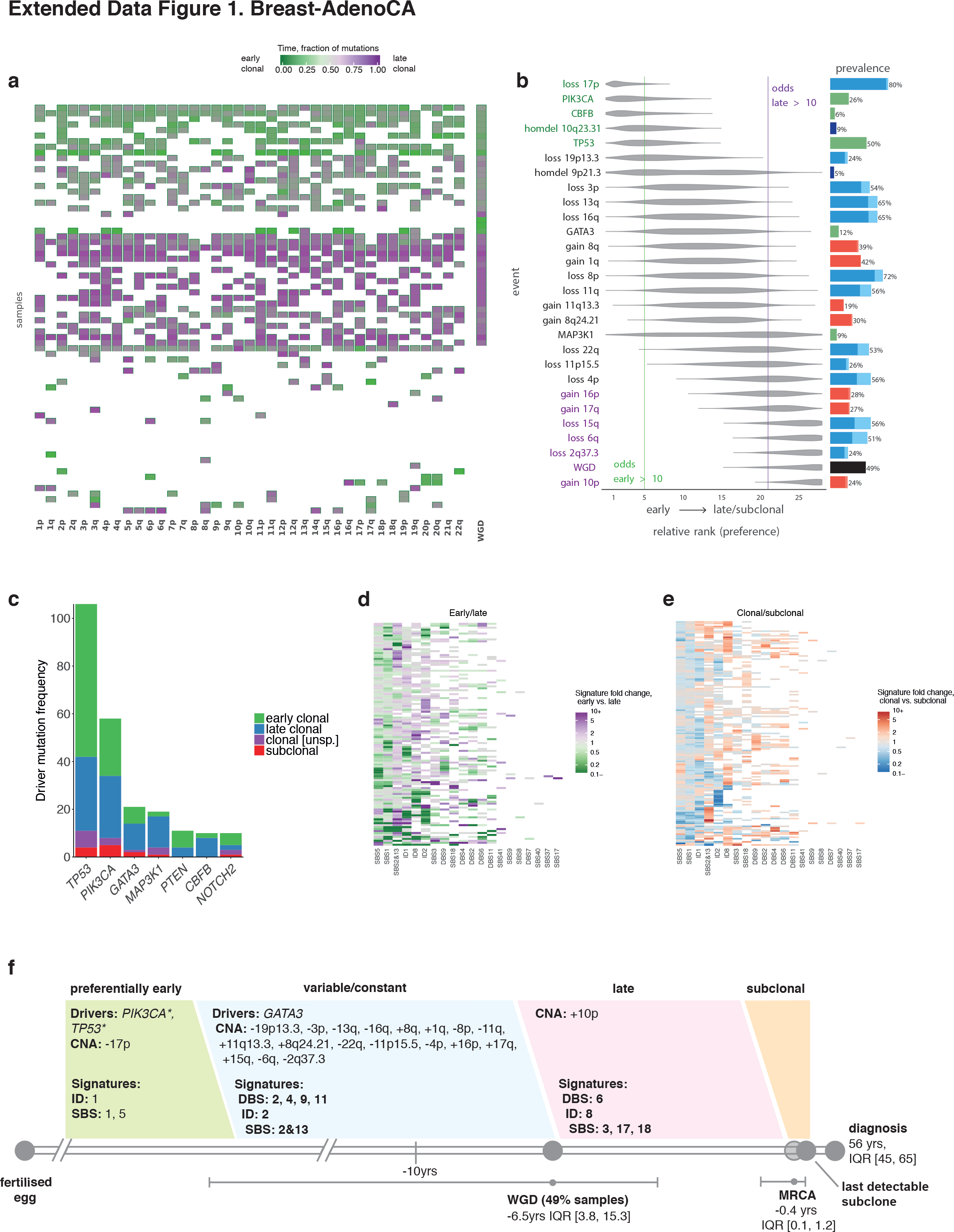
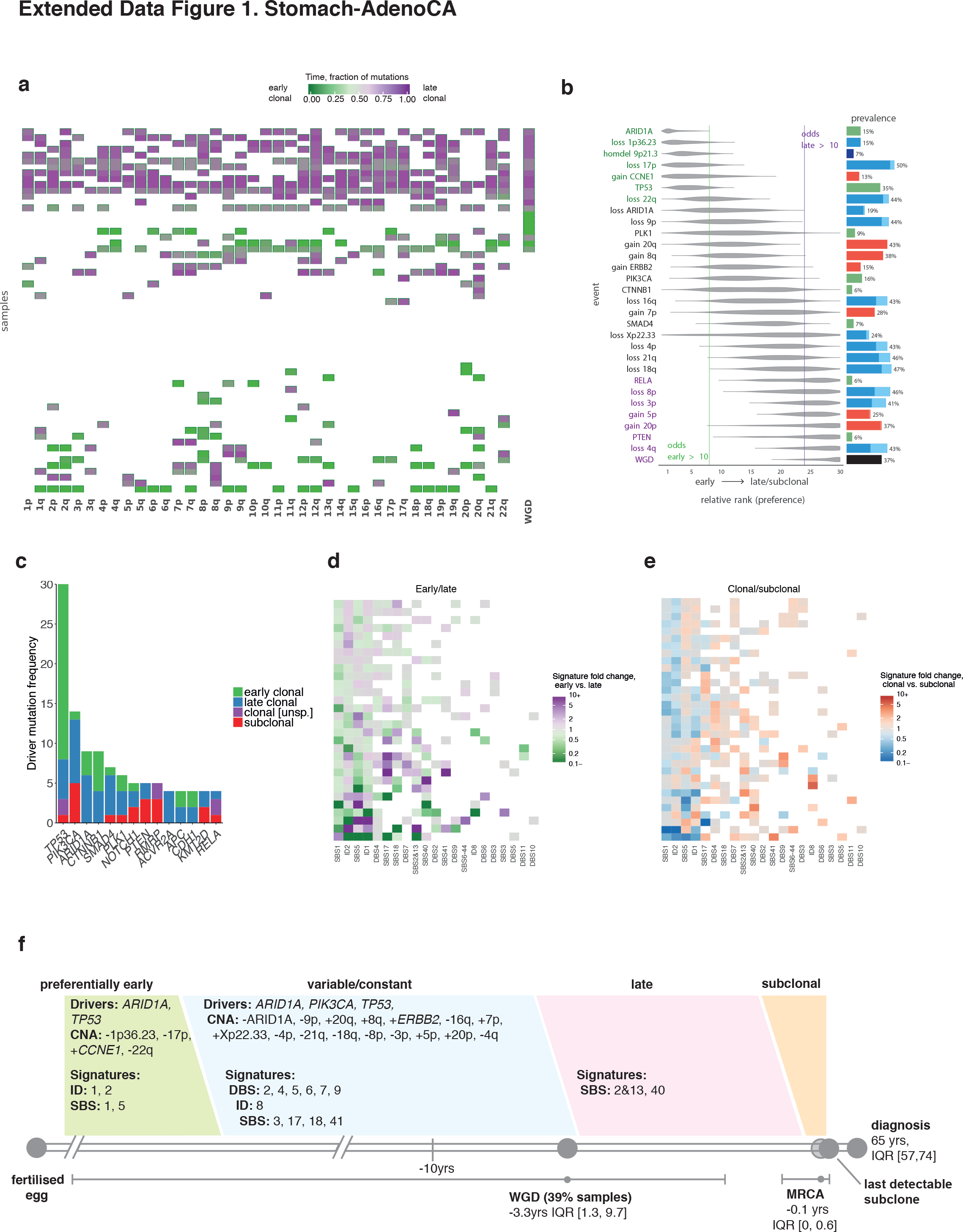
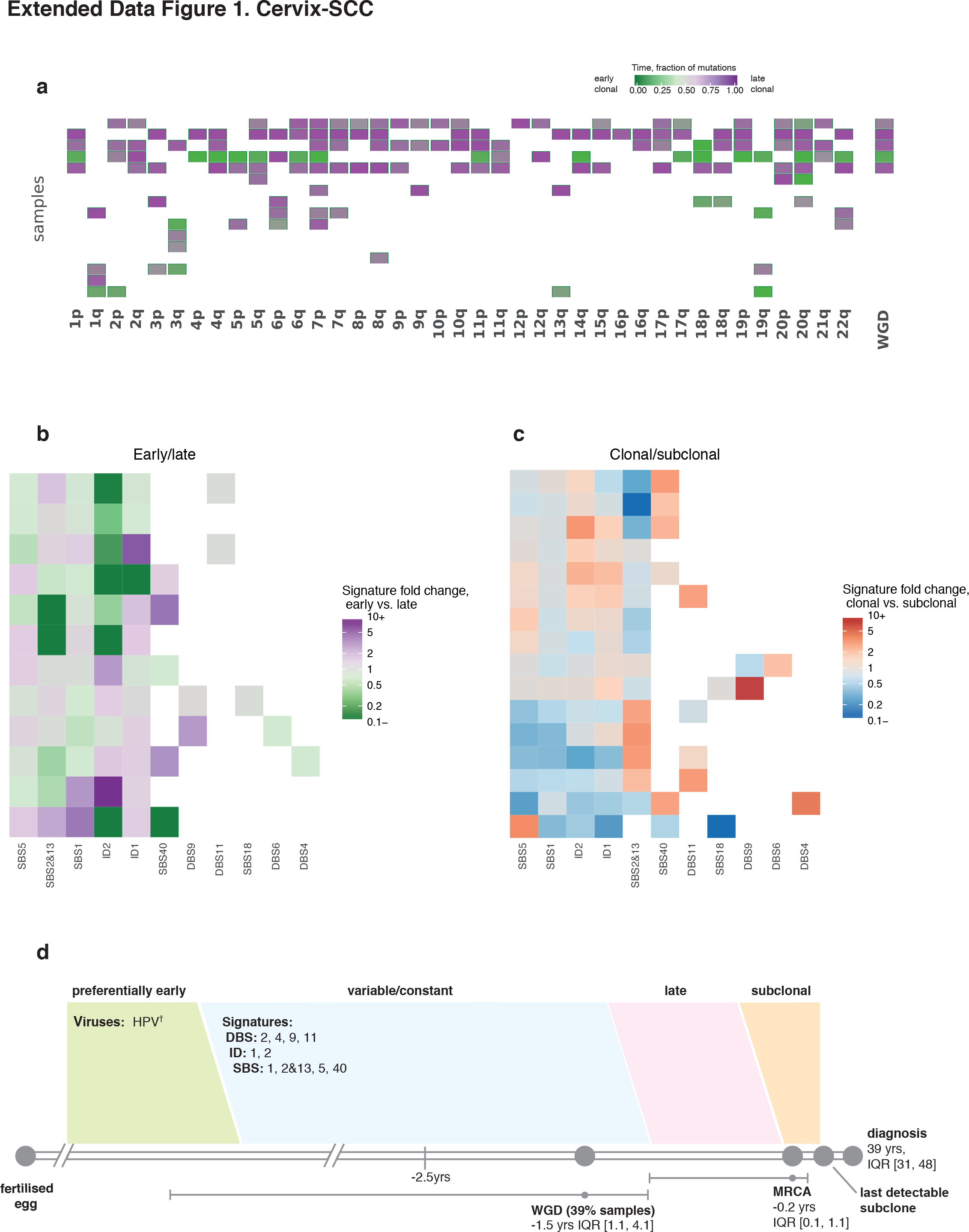
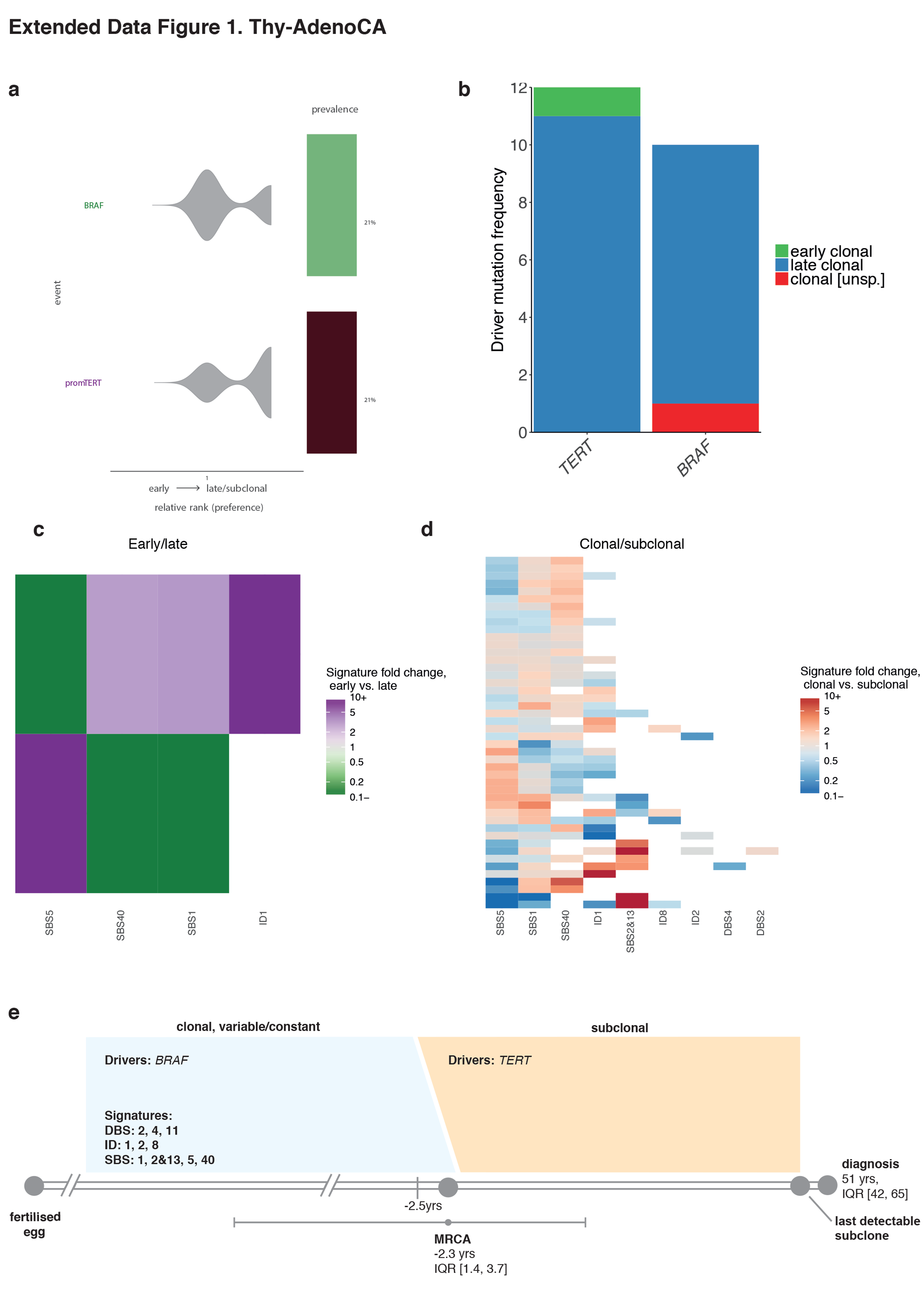
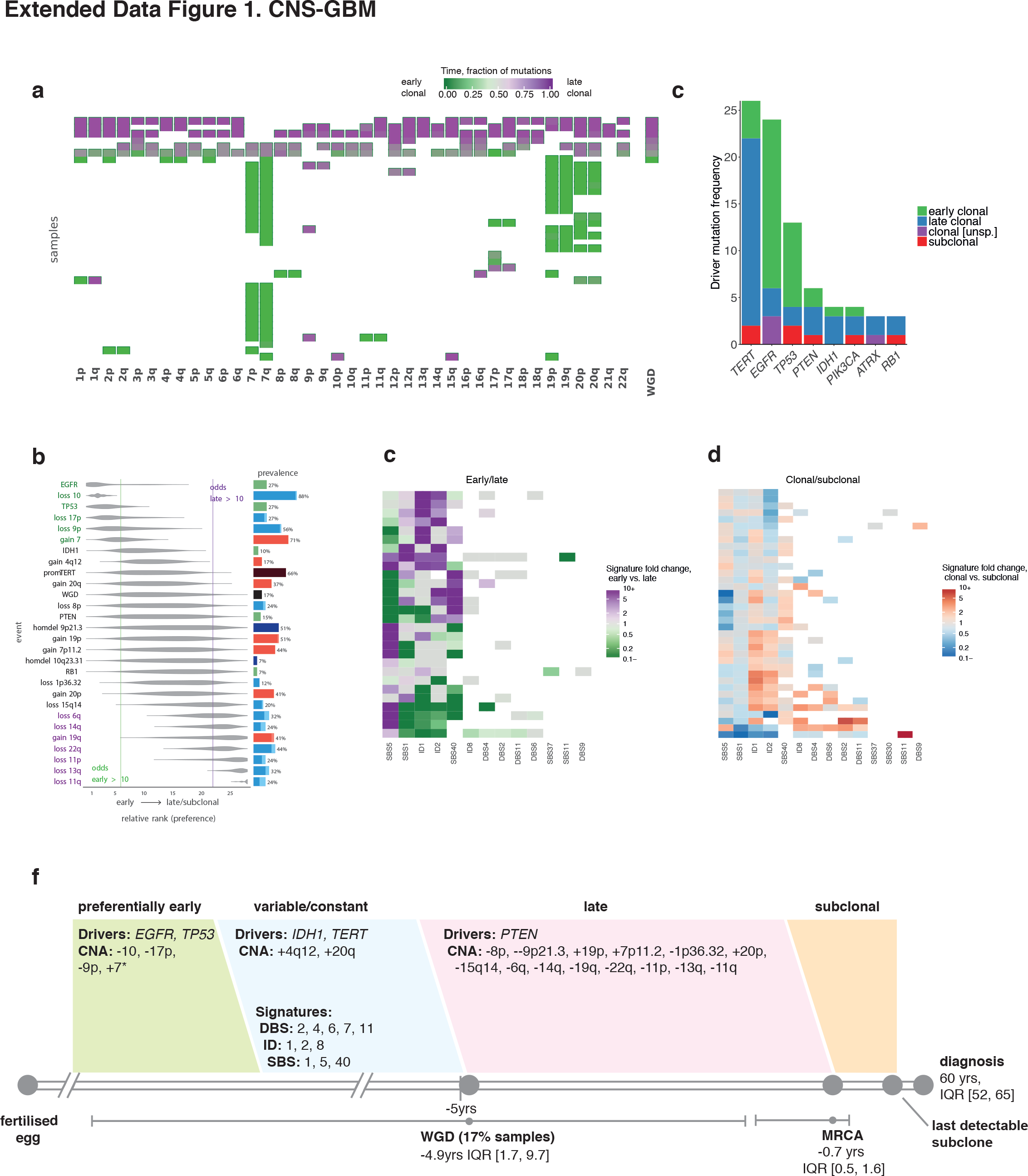
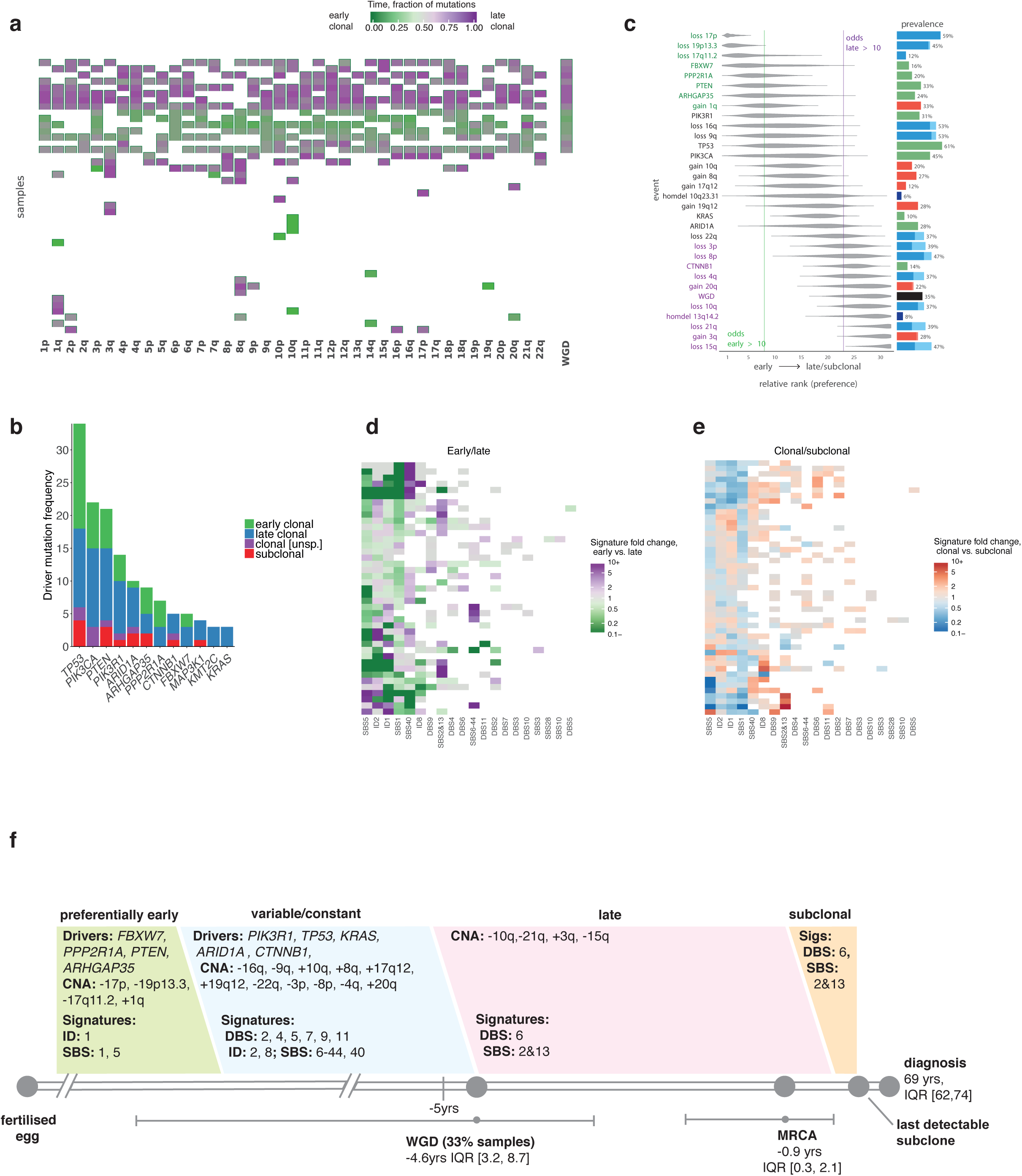
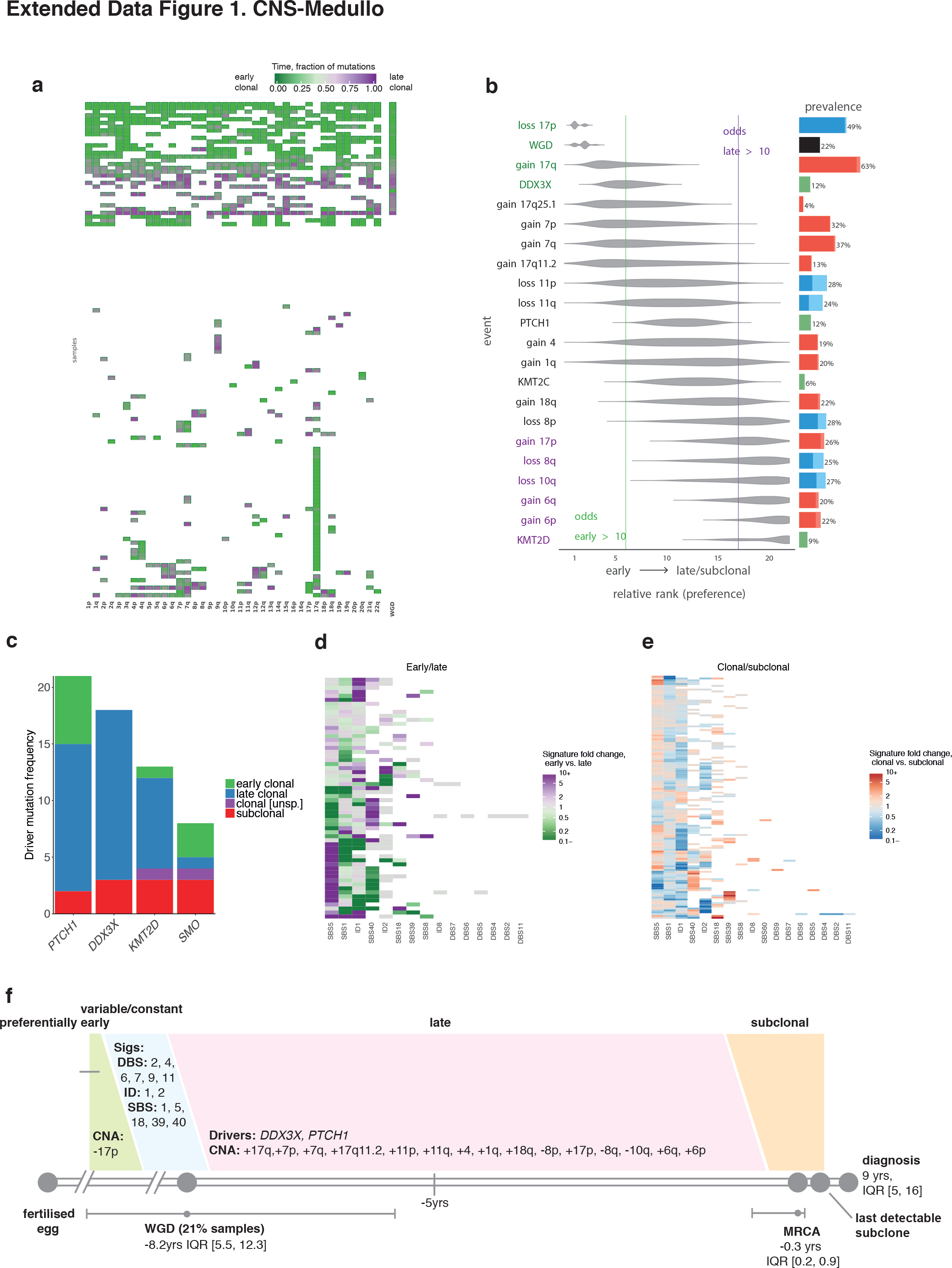
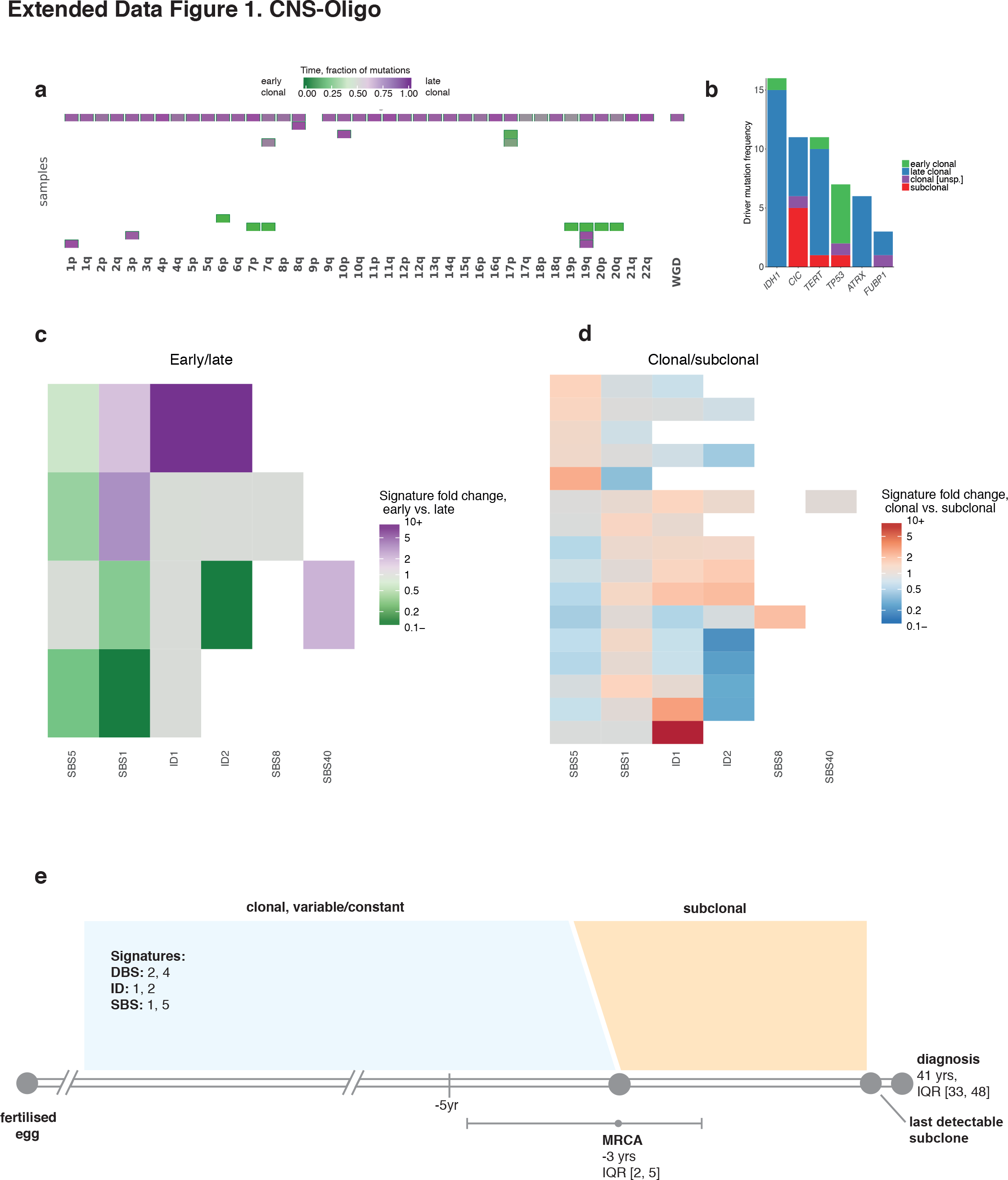
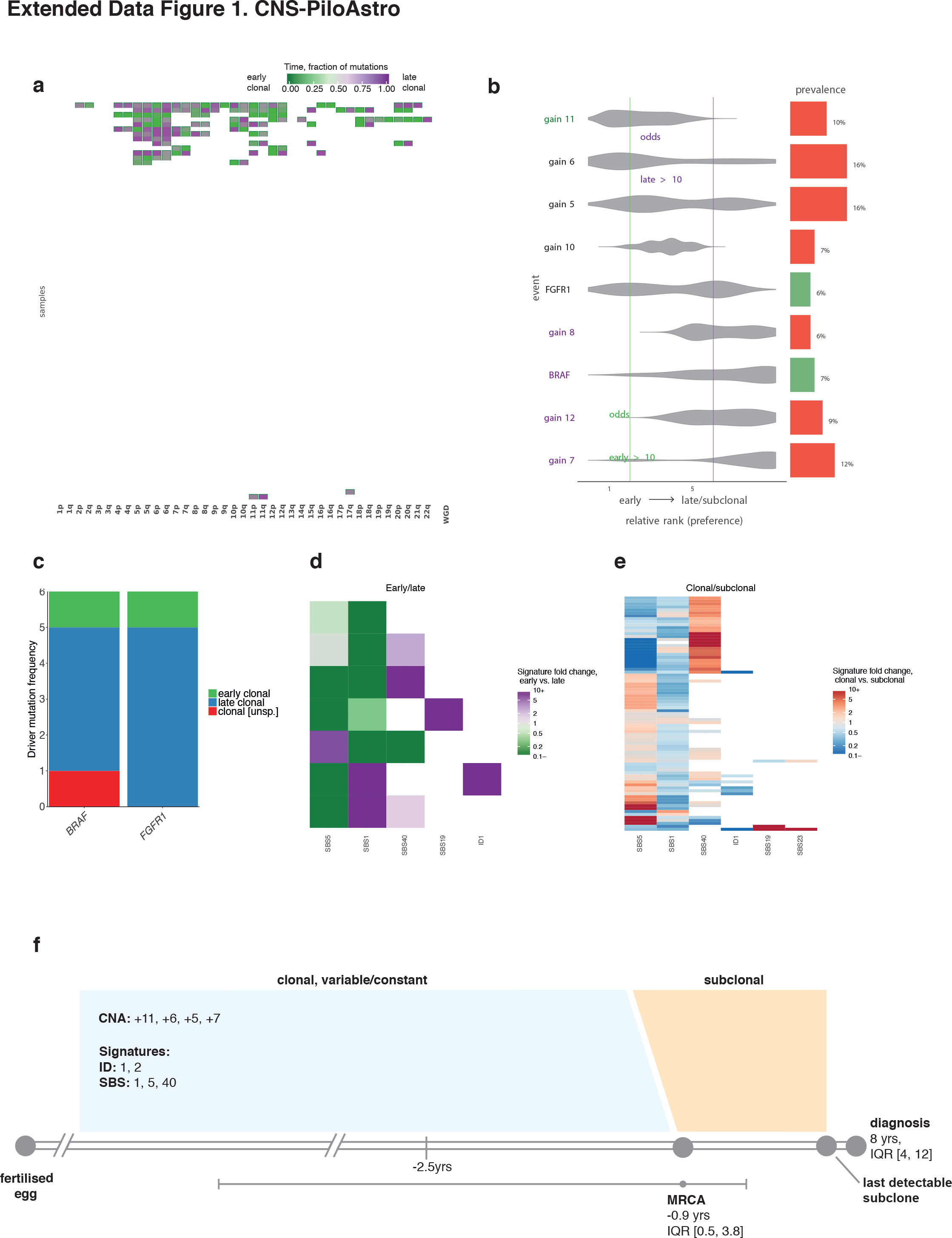
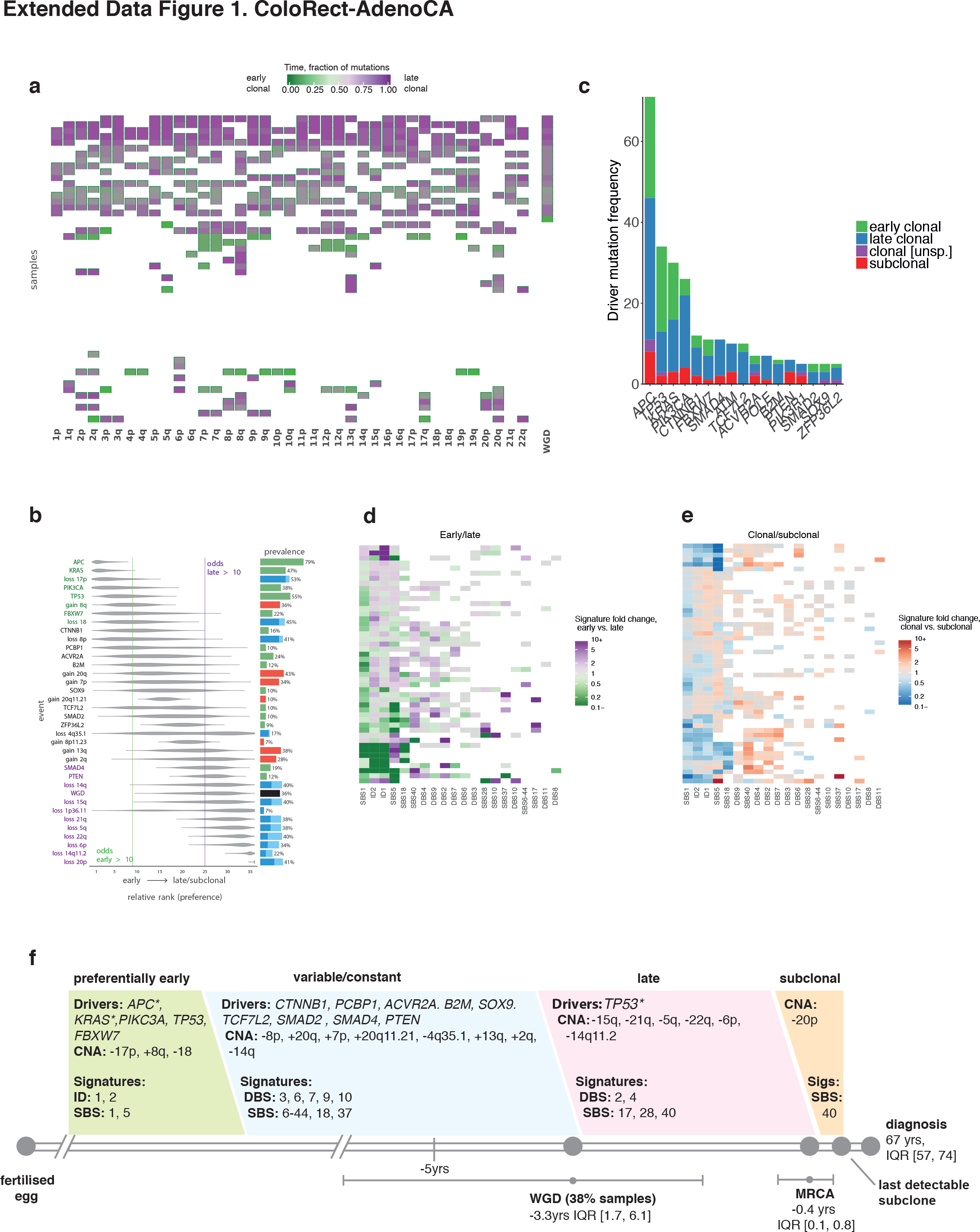
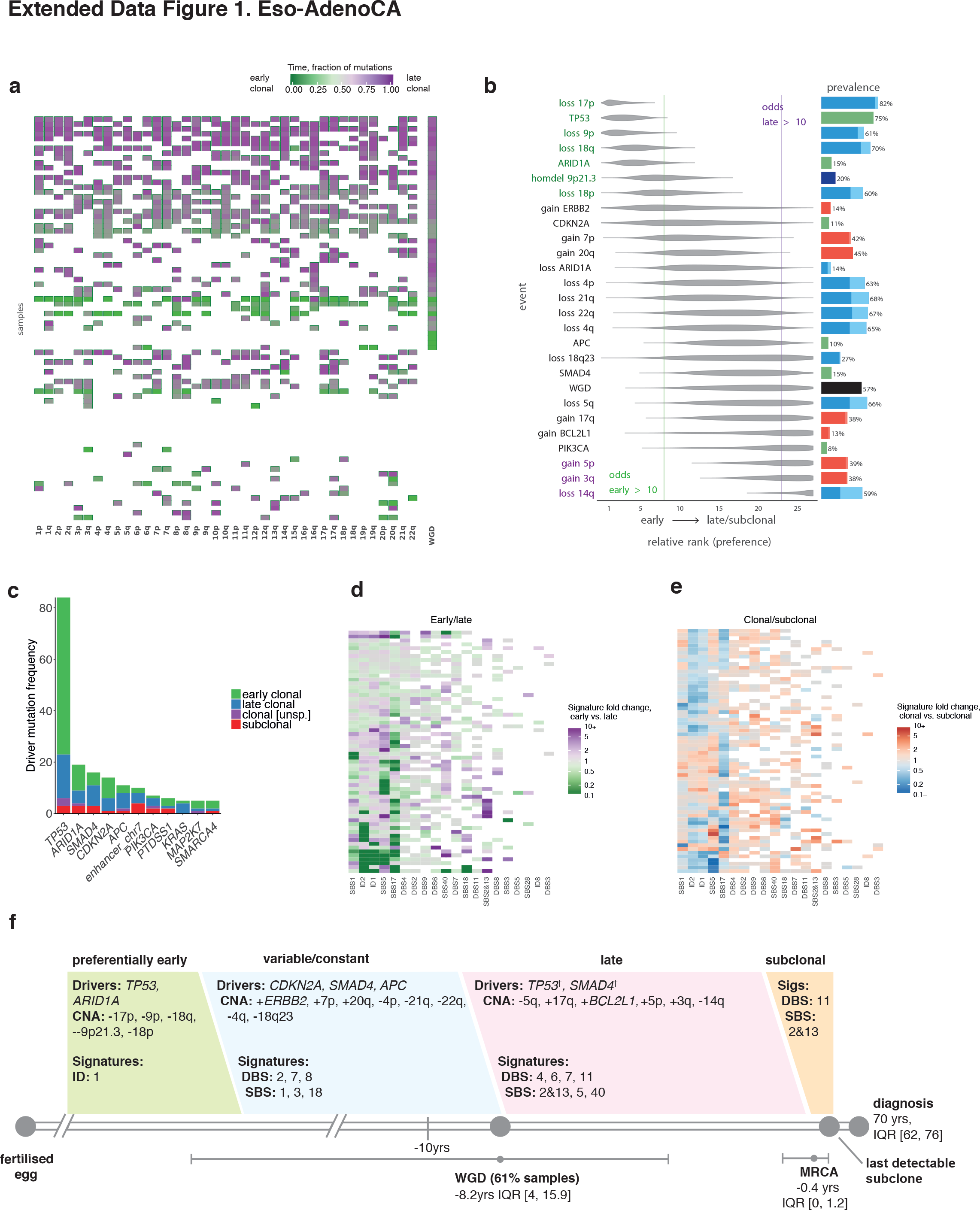
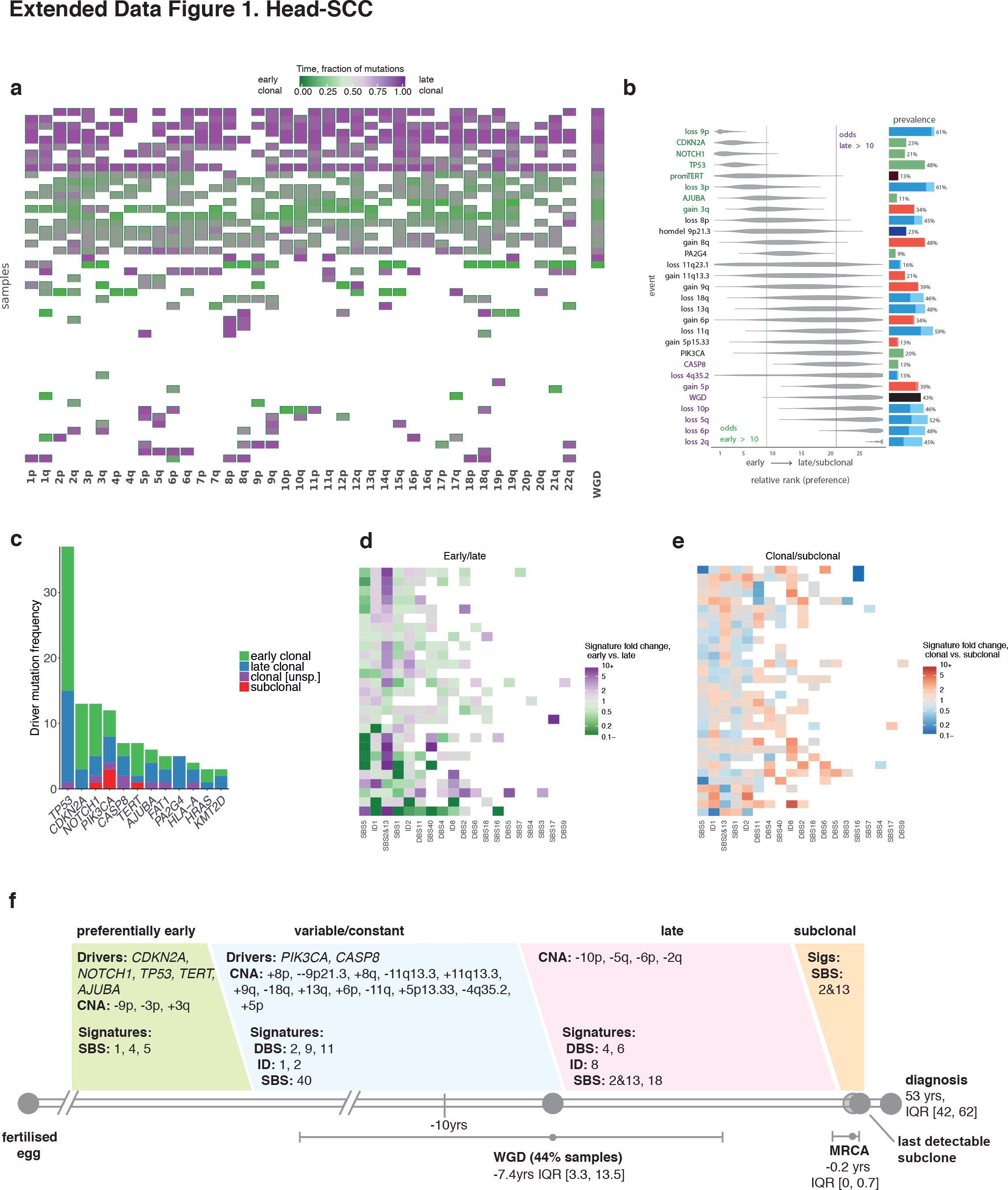
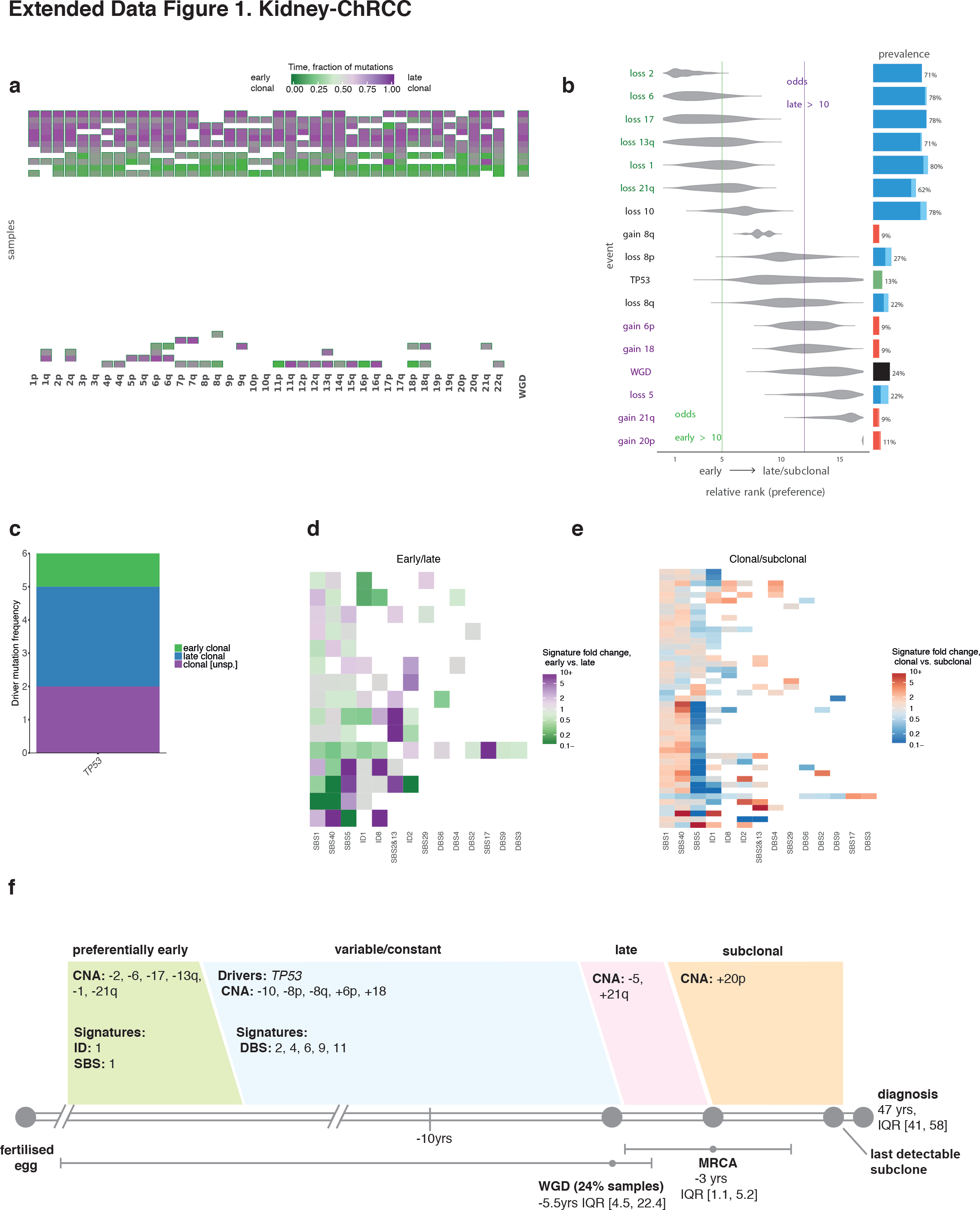
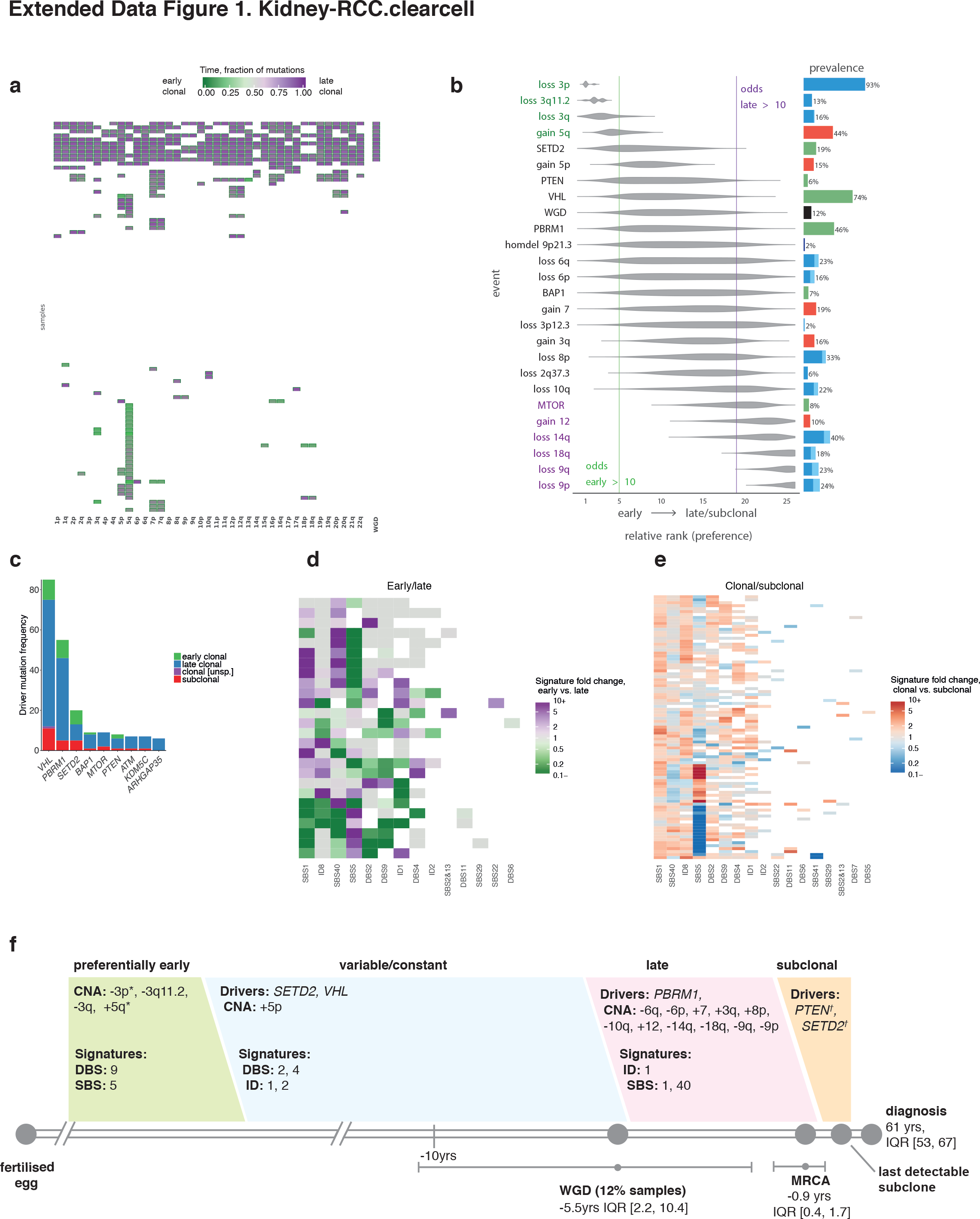
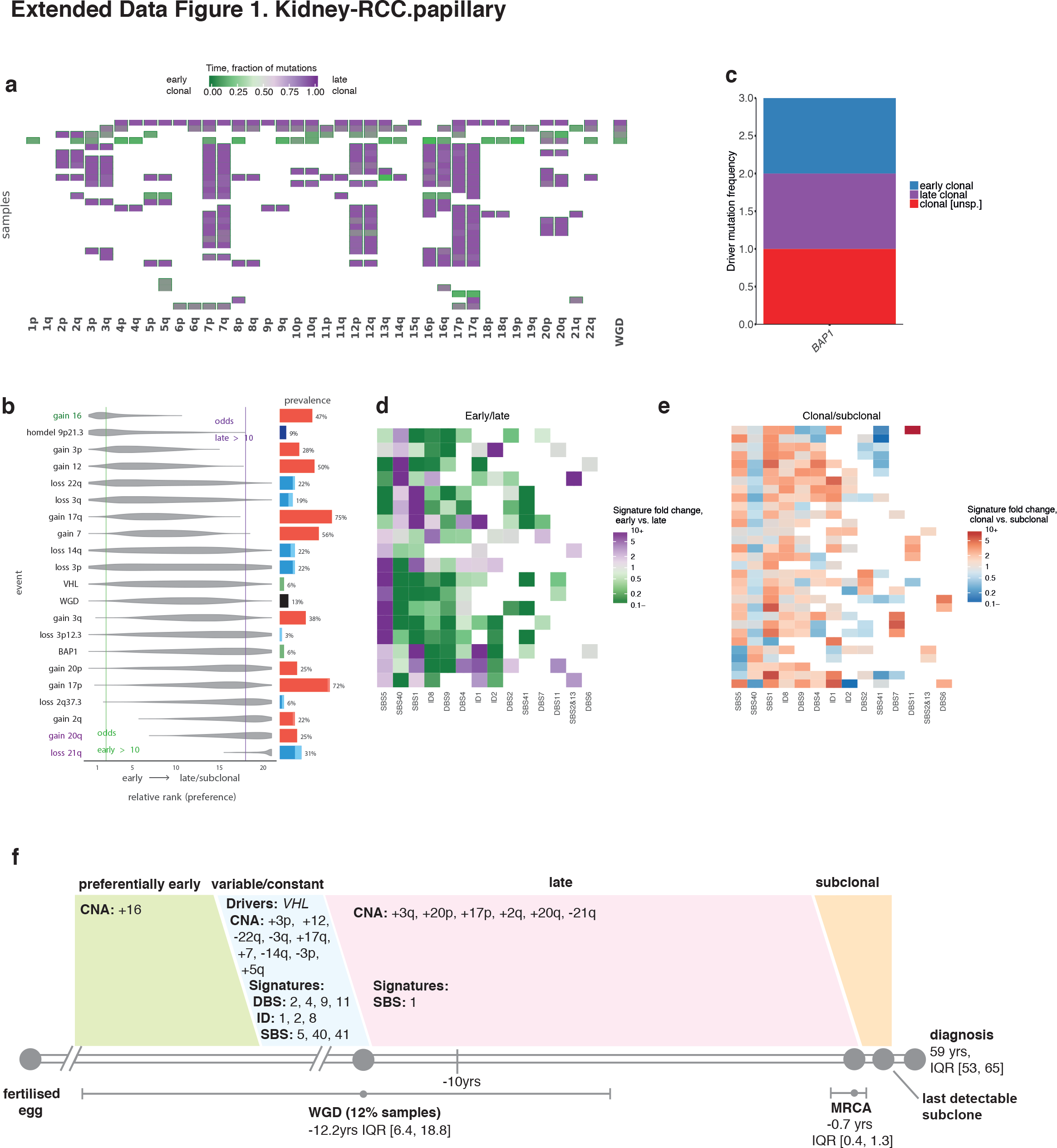
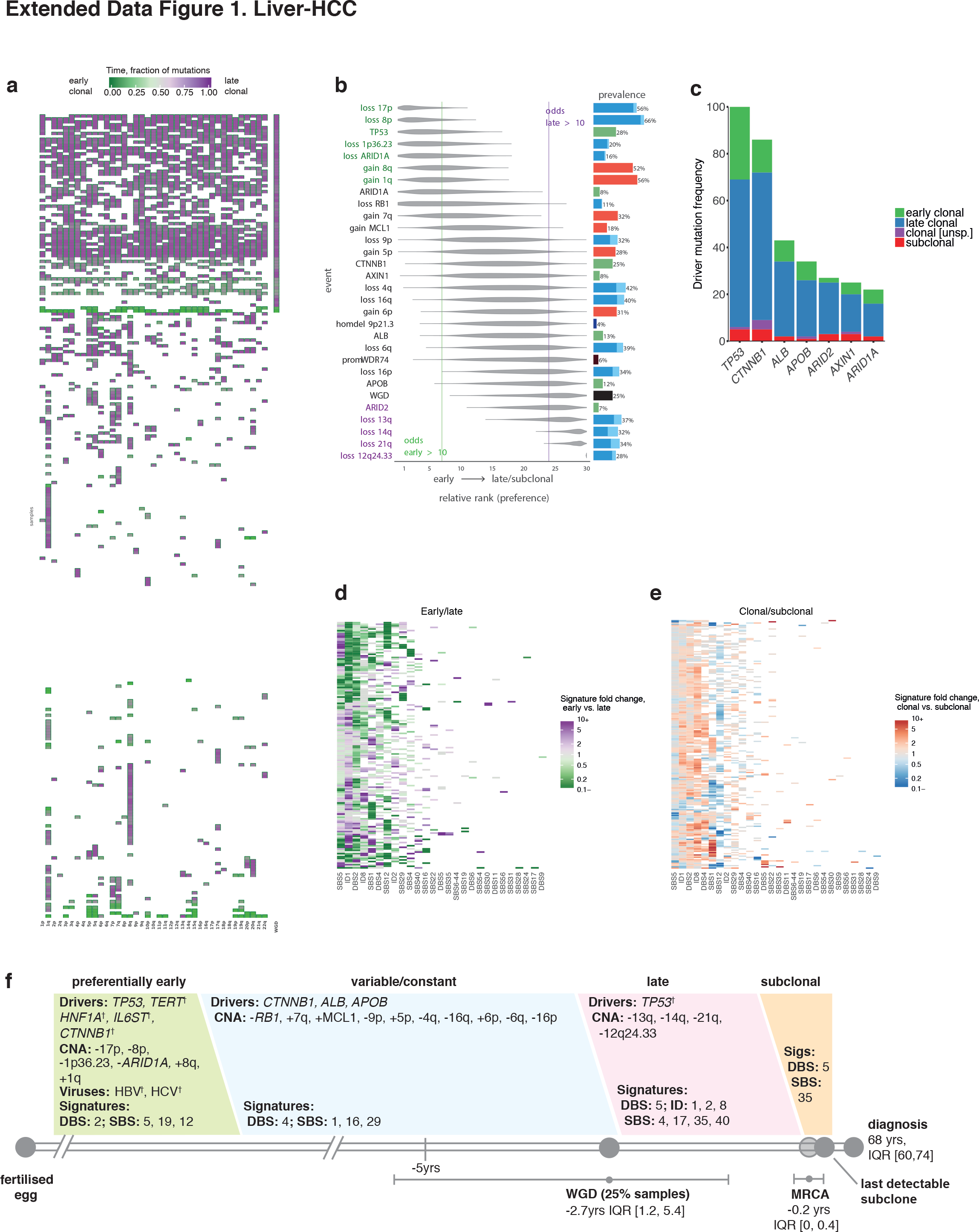
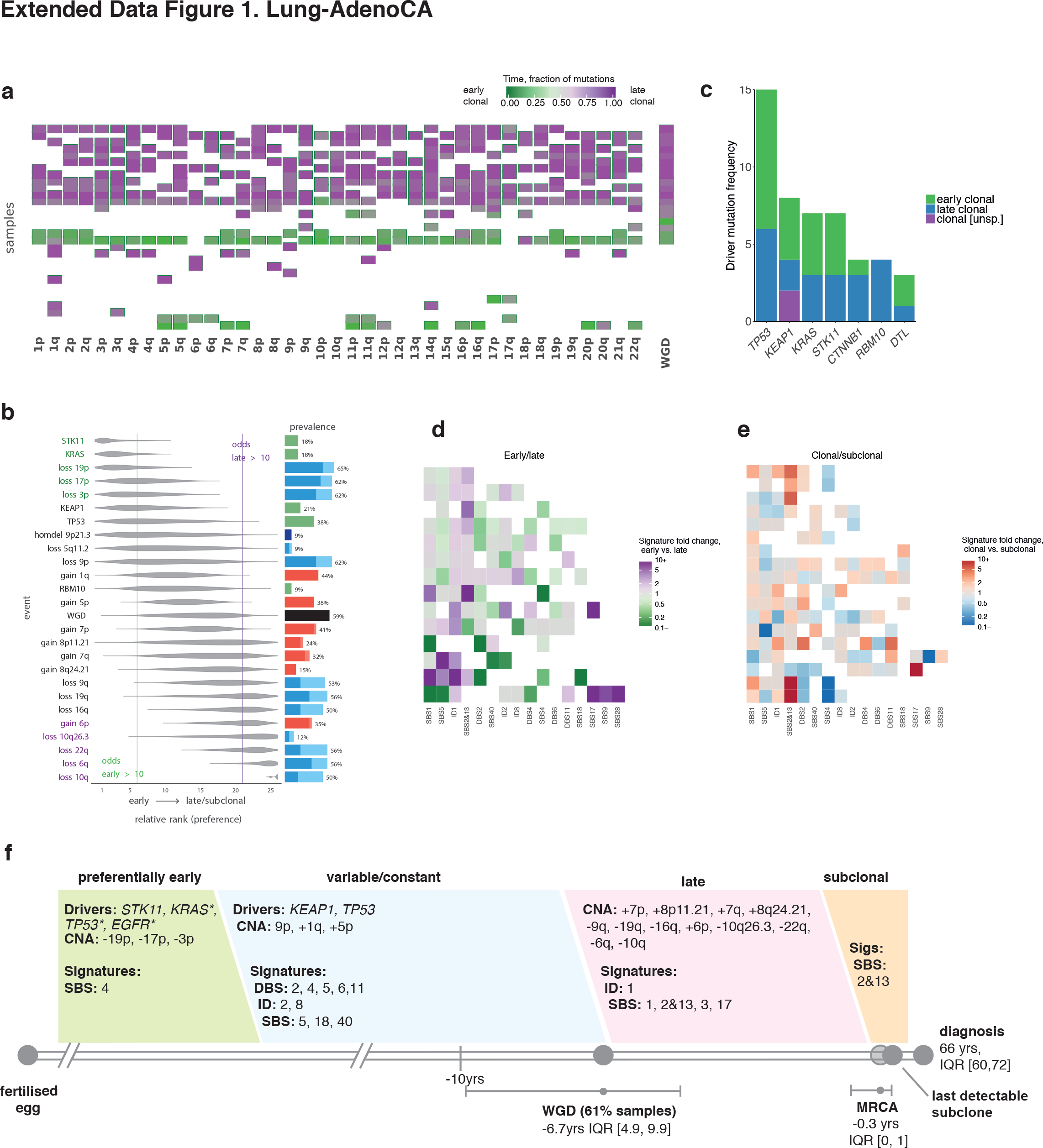
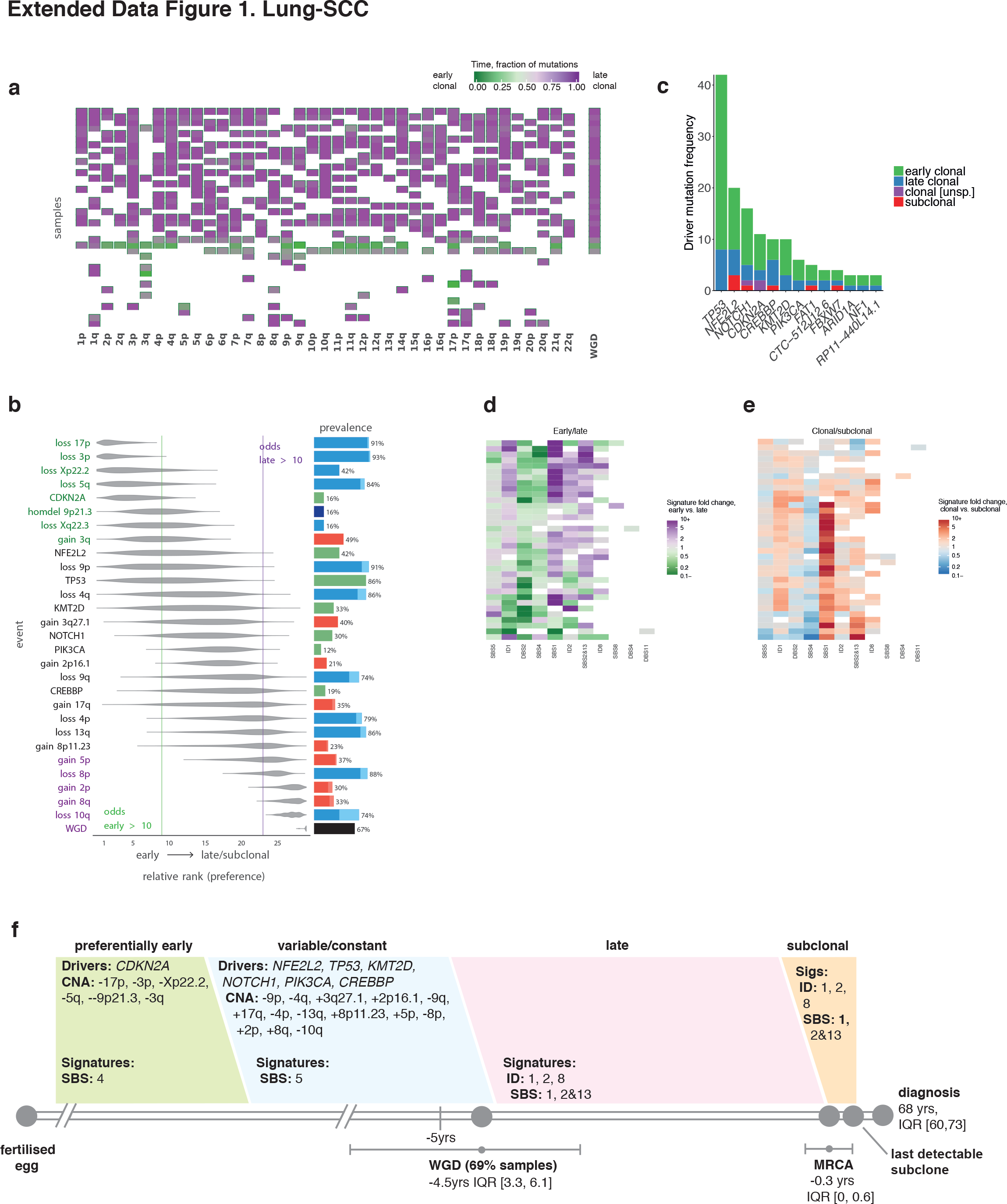
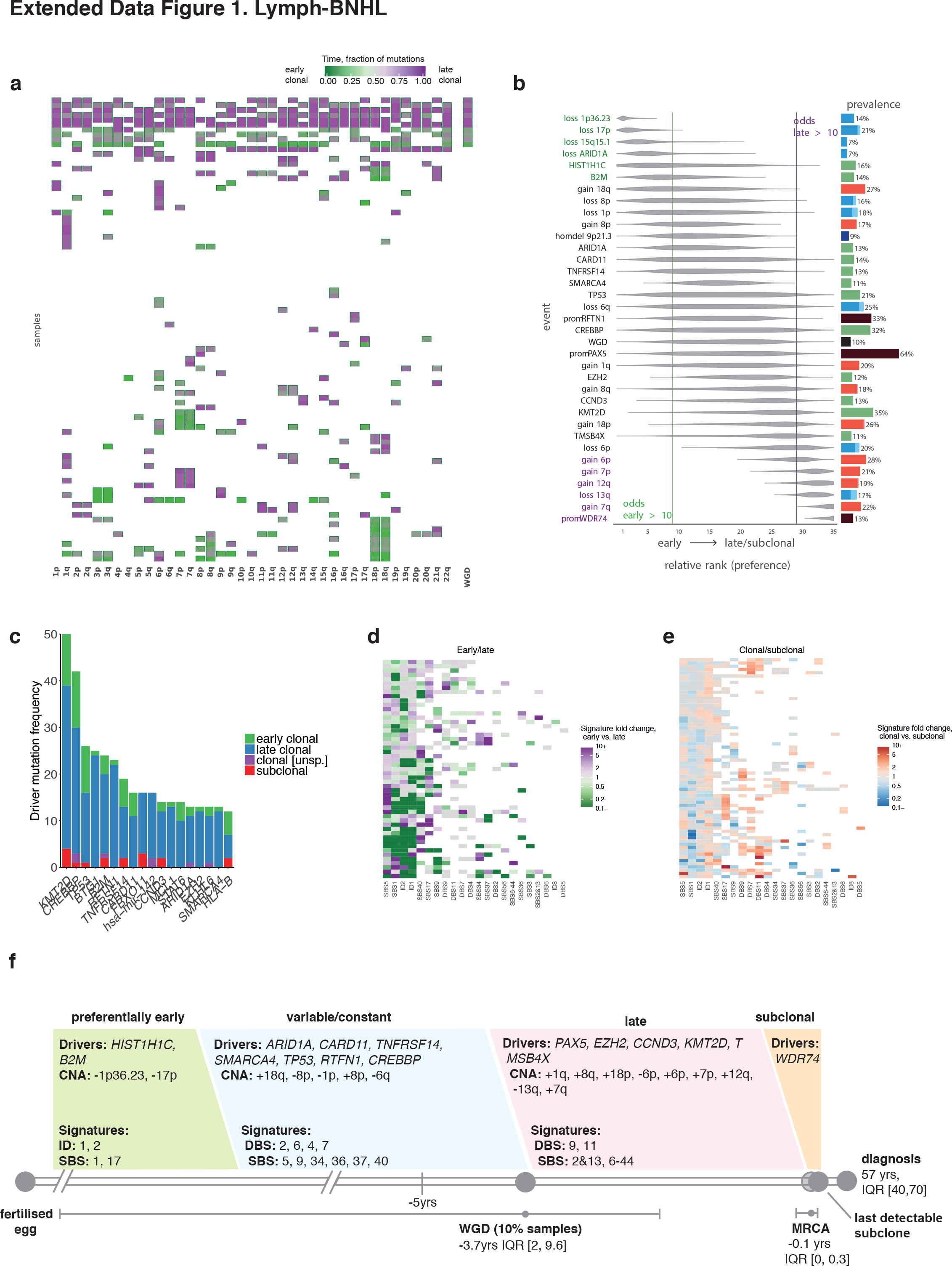
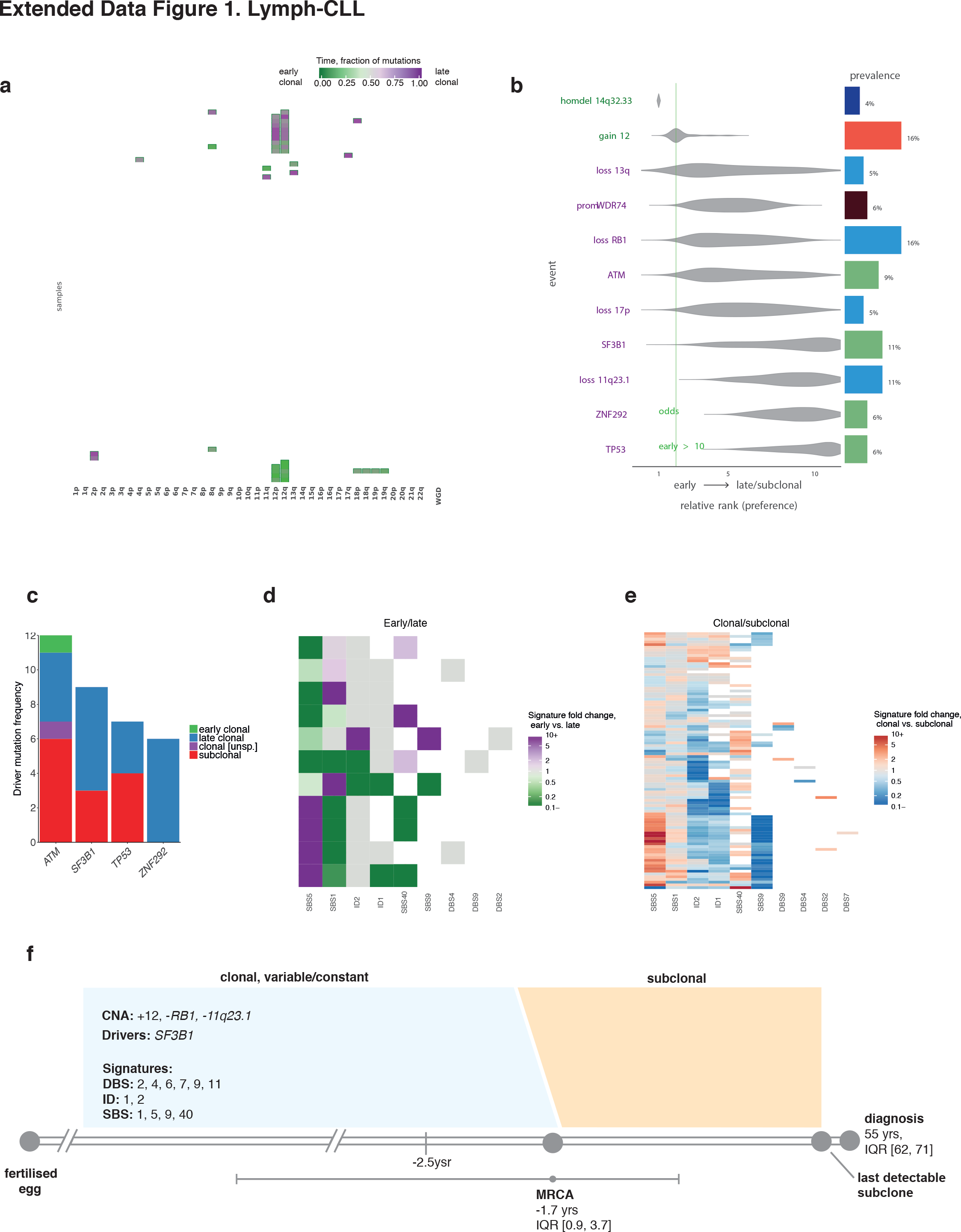
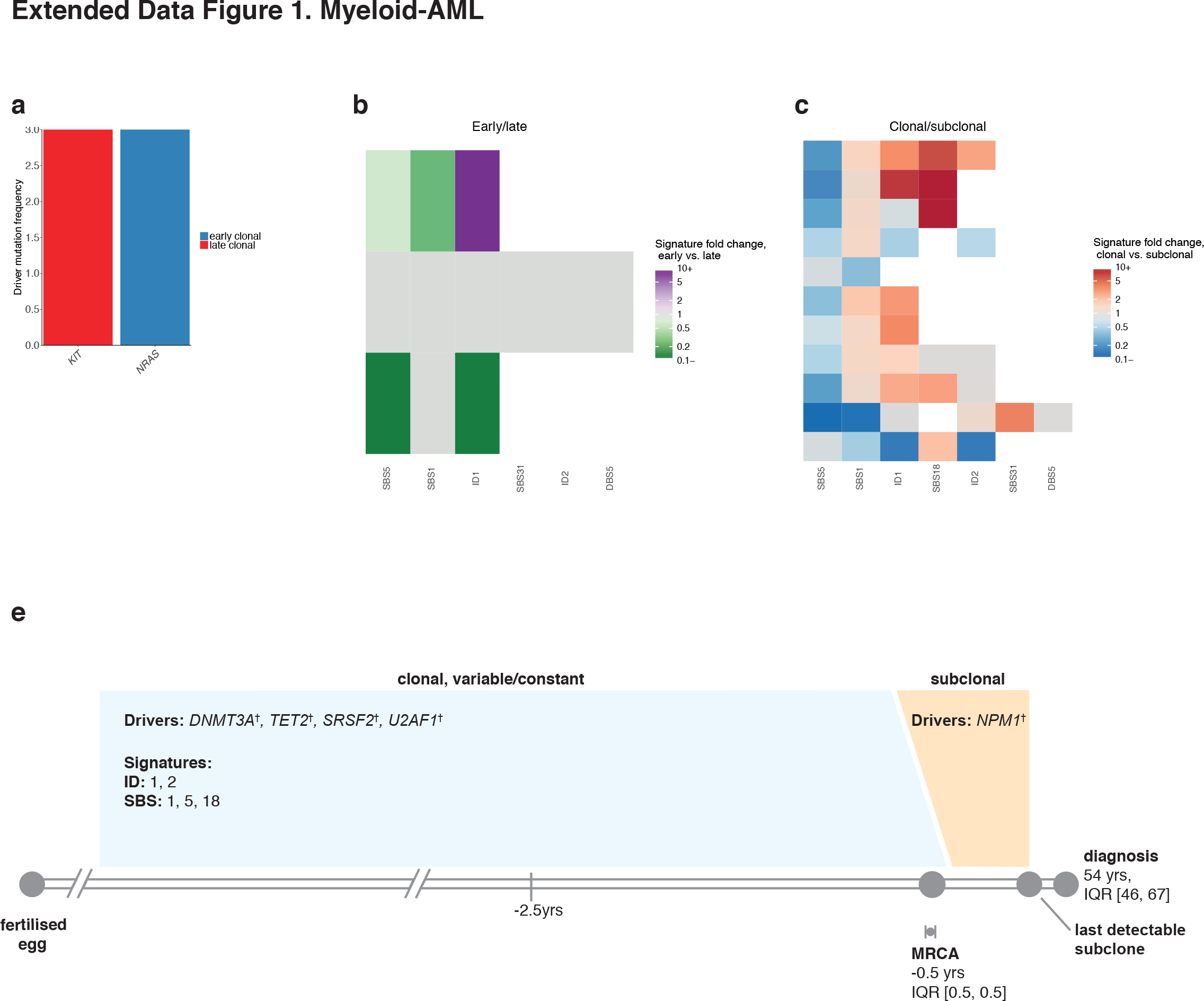
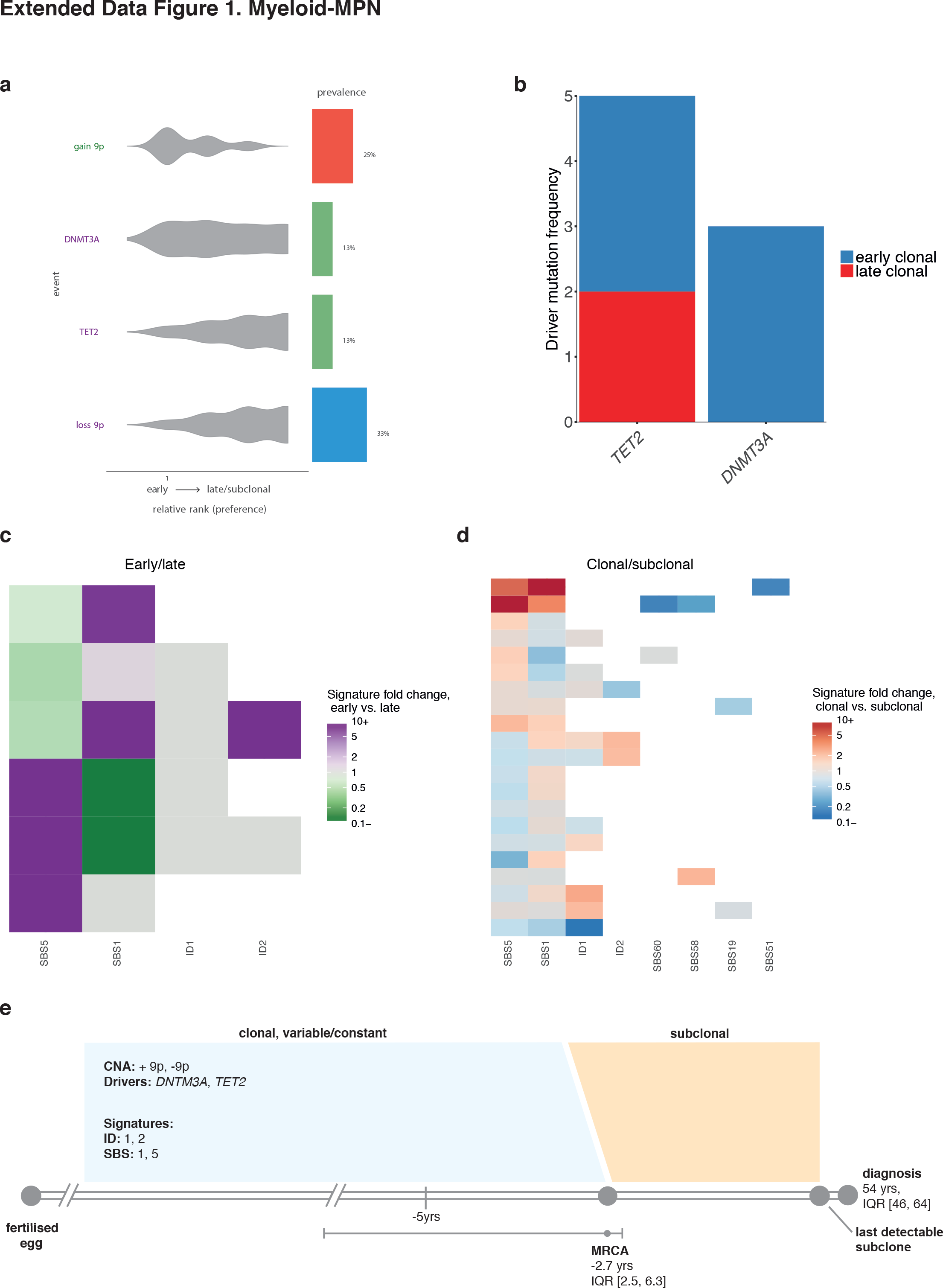
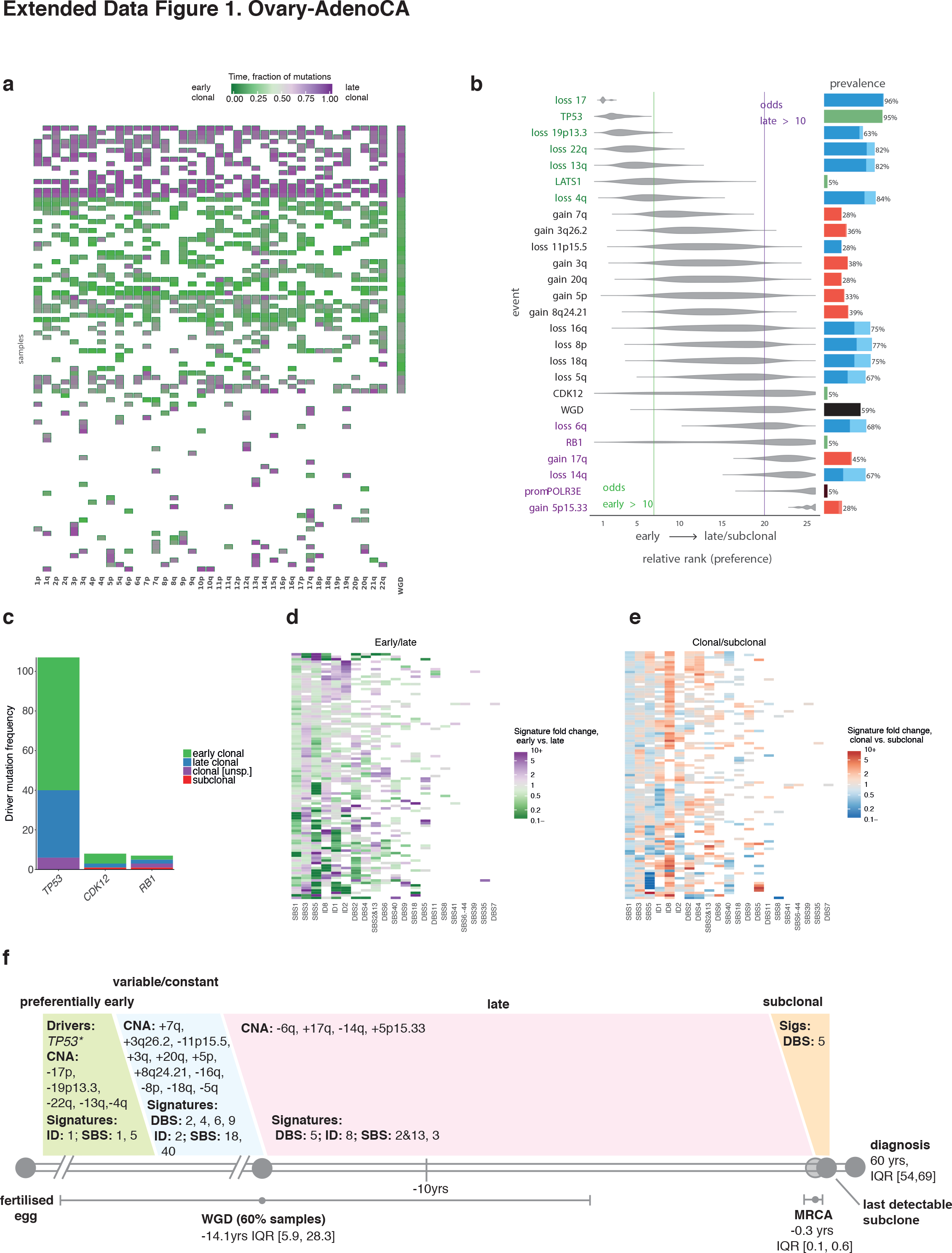
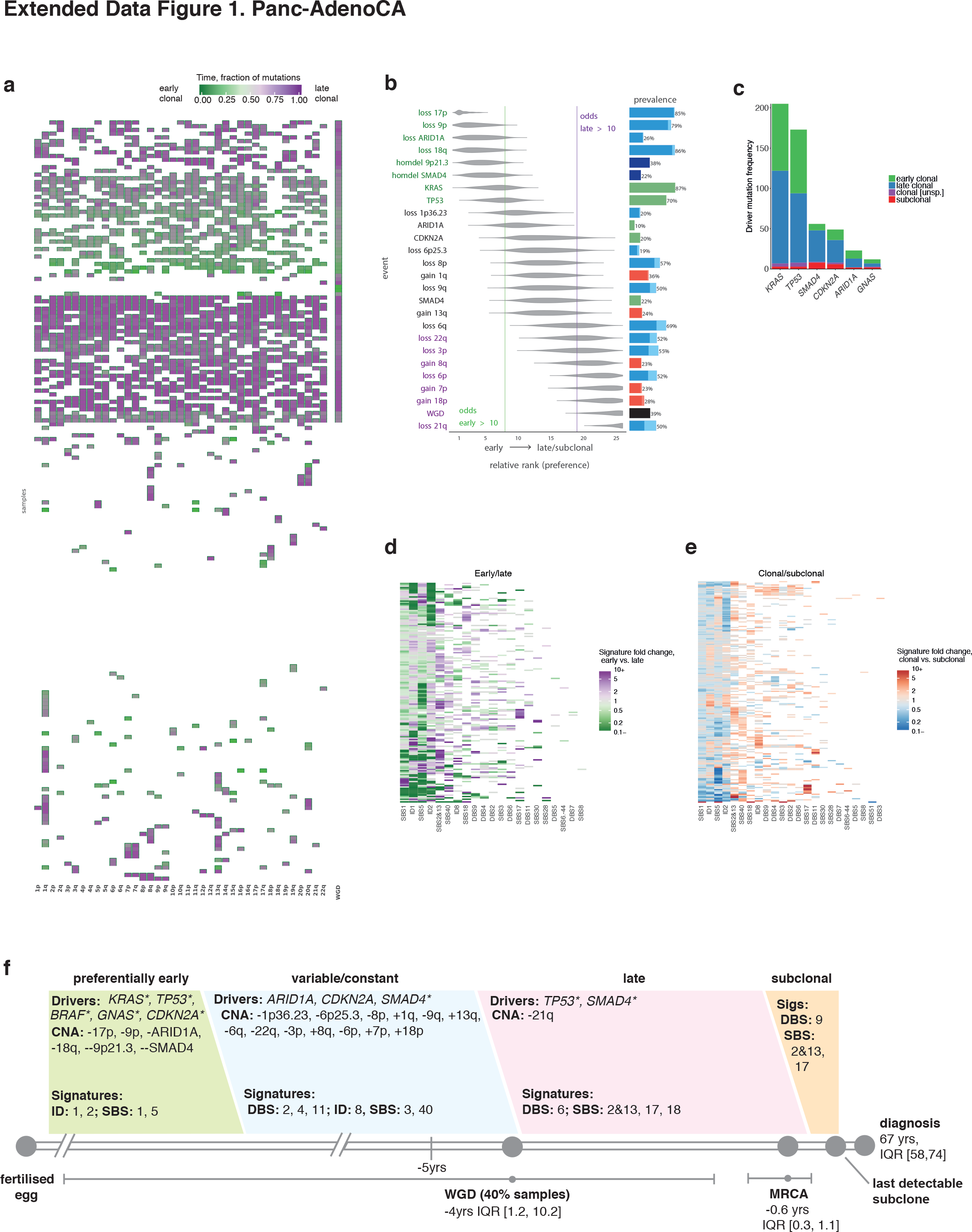
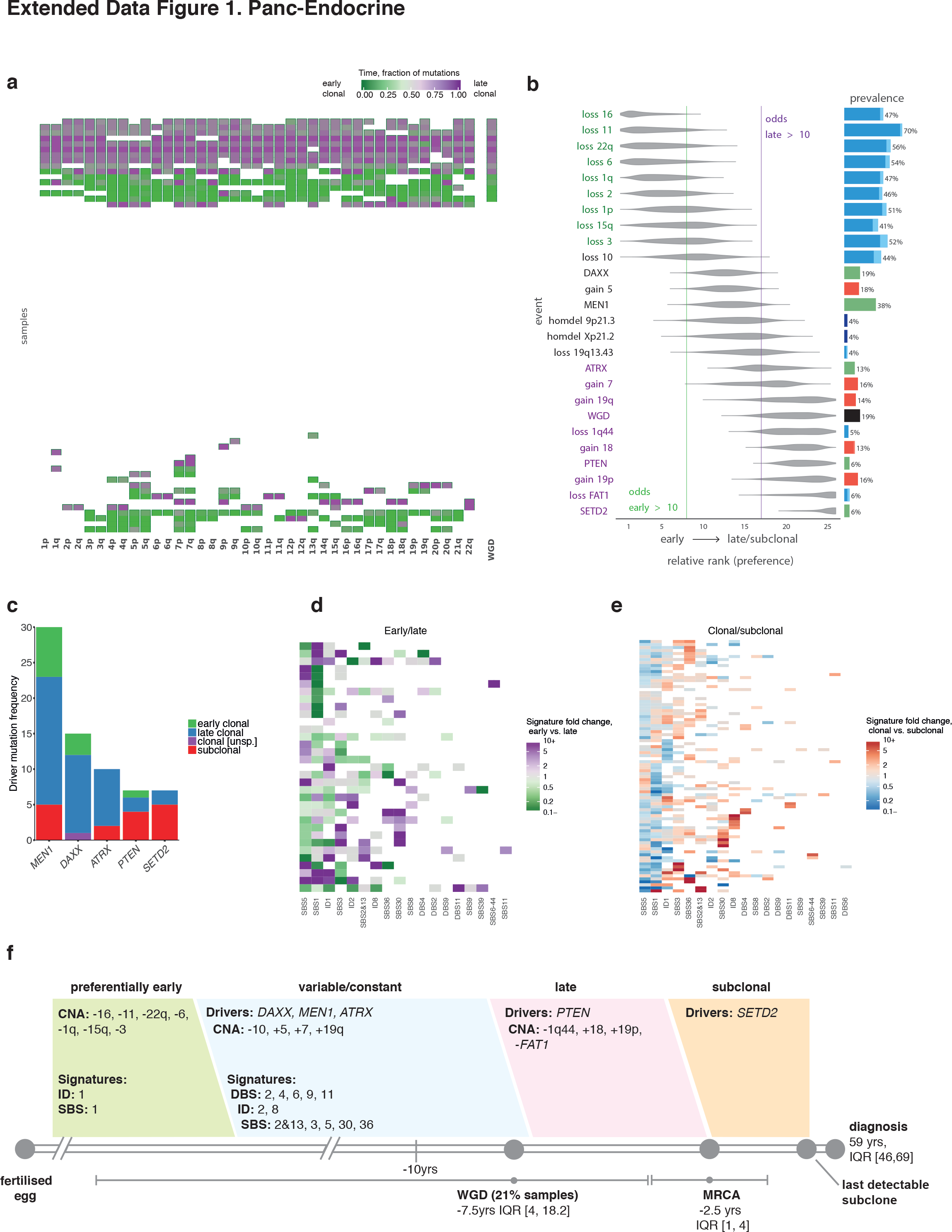
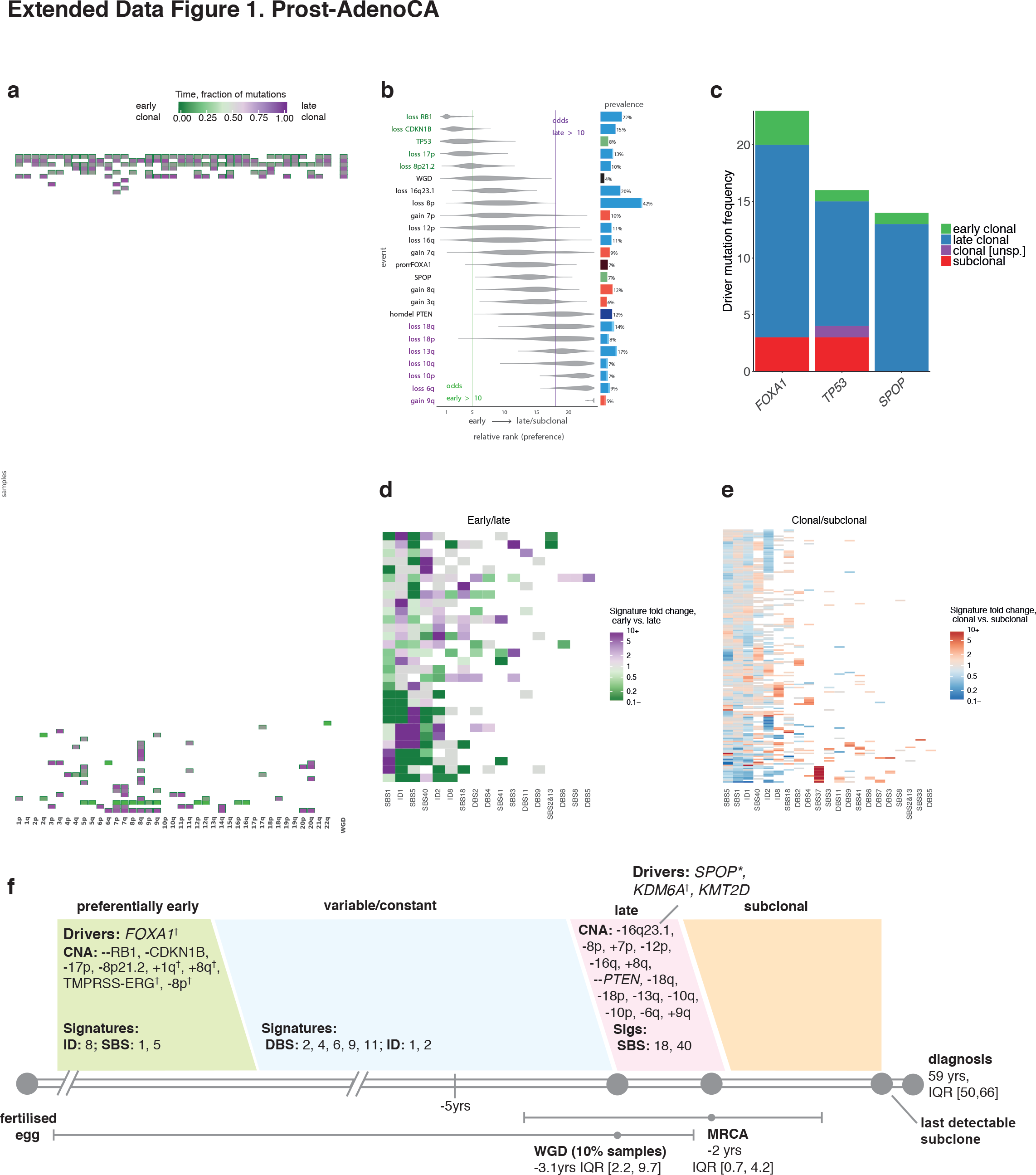
Summary of all results obtained per cancer type. (**a**) Clustered heatmaps of mutational timing estimates for gained segments, per patient. Colours as indicated in main text: green represents early clonal events, purple represents late clonal. (**b**) Relative ordering of copy number events and driver mutations across all samples per cancer type. (**c**) Distribution of mutations across early clonal, late clonal and subclonal stages, for the most common driver genes per cancer type. A maximum of 10 driver genes are shown. (**d**) Clustered mutational signature fold changes between early clonal and late clonal stages, per patient. Green and purple indicate, respectively, a signature decrease and increase in late clonal from early clonal mutations. Inactive signatures are coloured white. (**e**) As in (**d**) but for clonal versus subclonal stages. Blue indicates a signature decrease and red an increase in subclonal from clonal mutations. (**f**) Typical timeline of tumour development, per cancer type.

**Extended Data Figure 2.**
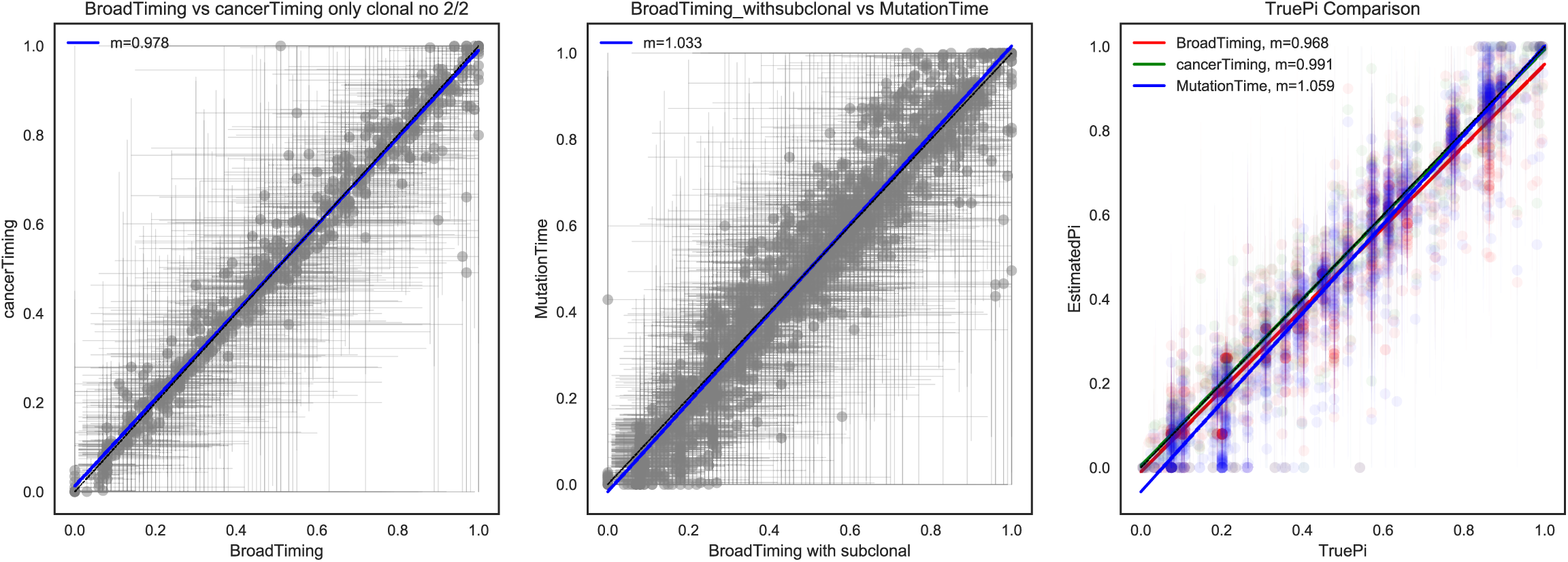
Comparison of methods used for timing of individual copy-number gains. (**a-b**) Pairwise comparison of the three approaches for timing individual copy number gains and (**c**) comparison with the simulated truth, showing high concordance.

**Extended Data Figure 3.**
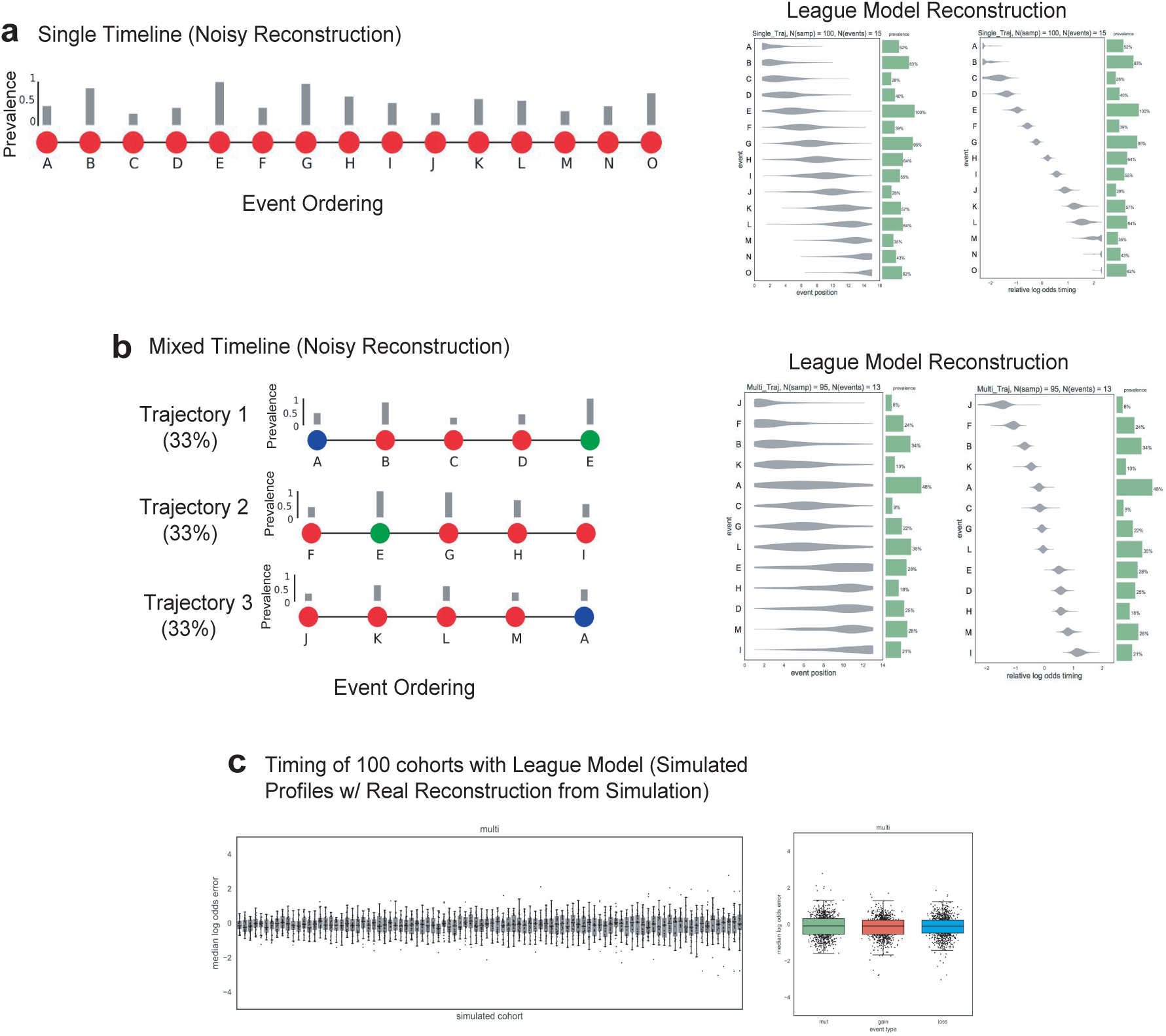
Validation of League model relative ordering reconstruction based on simulated cohorts of whole-genome samples. (**a**) League model results on a simulated cohort of samples from a single generalized relative order of events (with varied prevalence) showing high concordance with the true trajectory. (**b**) League model results on a simulated cohort of samples from a complex mixture of trajectories with different order of events showing high concordance with the expected average trajectory. (**c**) Estimation of accuracy of the League model reconstruction by simulation of a set of 100 cohorts with random trajectory mixtures and quantifying the distance in log odds early/late from perfect ordering. For the vast majority of events (even with low number of occurrences in the cohort) the log odds error does not exceed 1, confirming that barely any of the events would switch between timing categories.

**Extended Data Figure 4.**
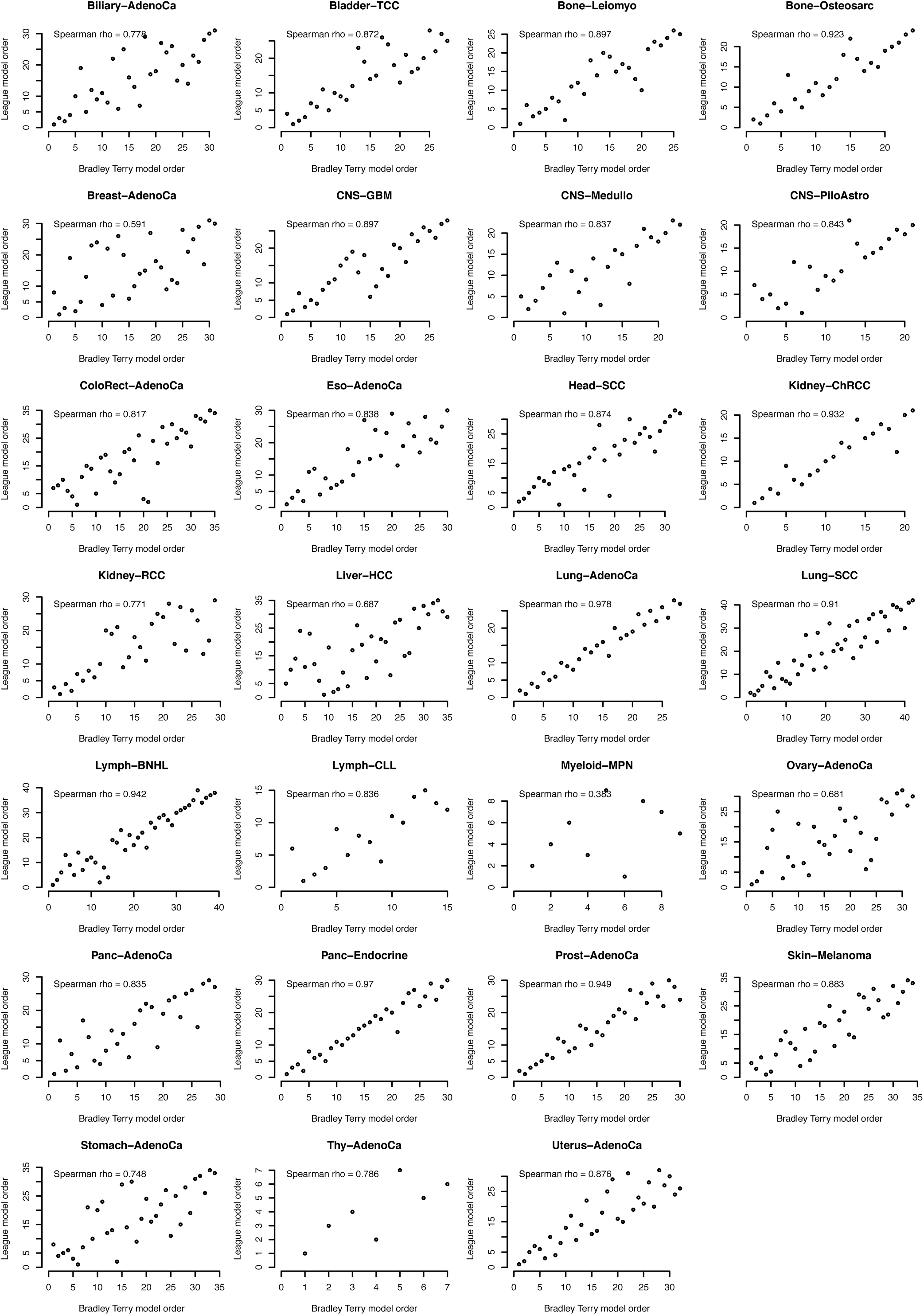
Correlation between League model and Bradley-Terry model ordering. Direct comparison for each tumour type of the League and Bradley-Terry models for determining the order of recurrent somatic mutations and copy number events. Axes indicate the ordered events observed in the respective tumour types. Correlation is quantified by Spearman’s rank correlation coefficient.

**Extended Data Figure 5.**
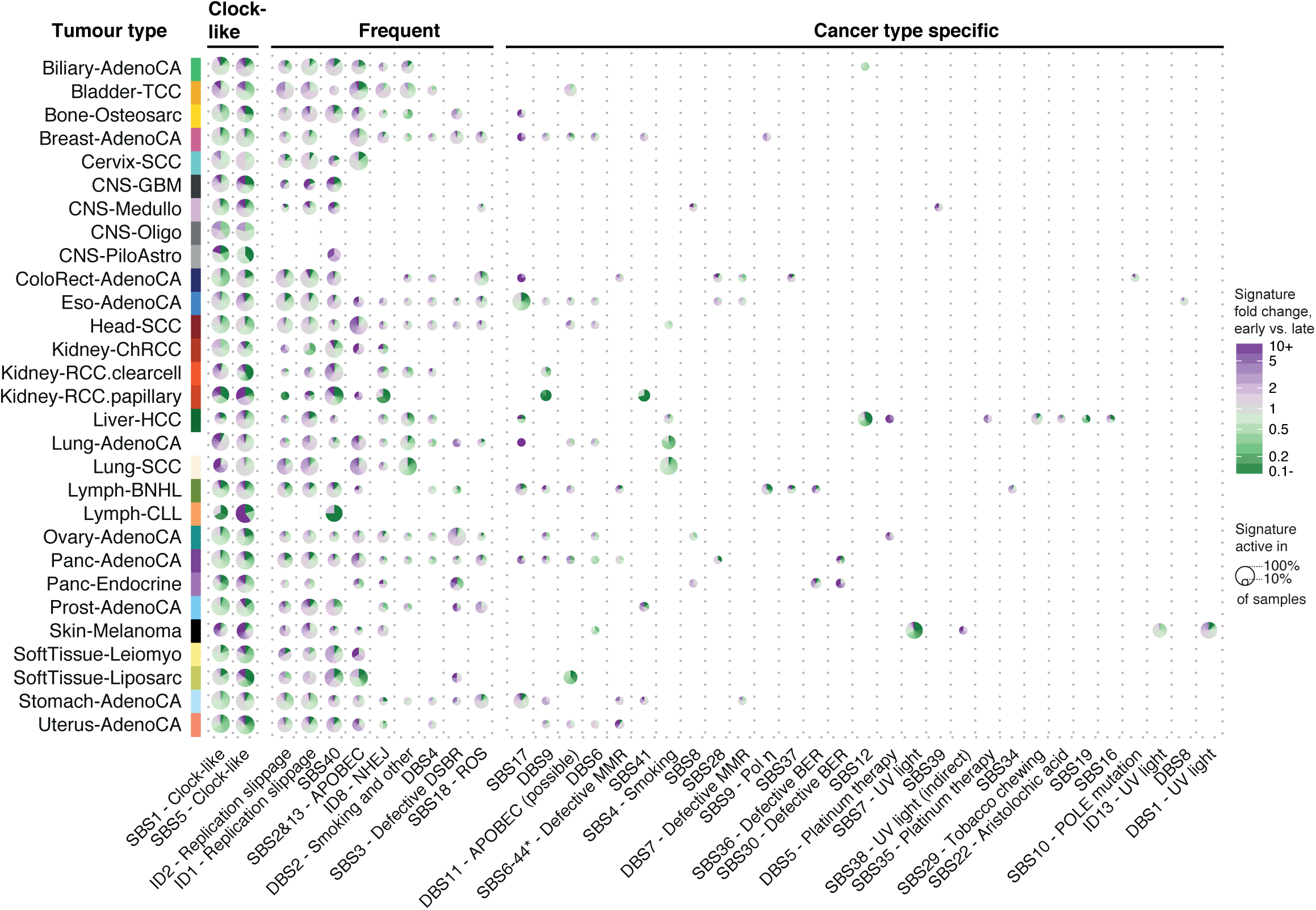
Overview of early to late clonal signature changes across tumour types. Pie charts representing signature changes per cancer type. The colour scheme is as above; signatures that decrease between early and late are coloured green, signatures that increase are purple. The size of each pie chart represents the frequency of each signature. Signatures are split into three categories: “clock-like”, comprising the putative clock signatures 1 and 5, “frequent”, which are signatures present in 10 or more cancer types, and “cancer type specific”, which are in fewer than 10 cancer types and are often limited to specific cohorts.

**Extended Data Figure 6.**
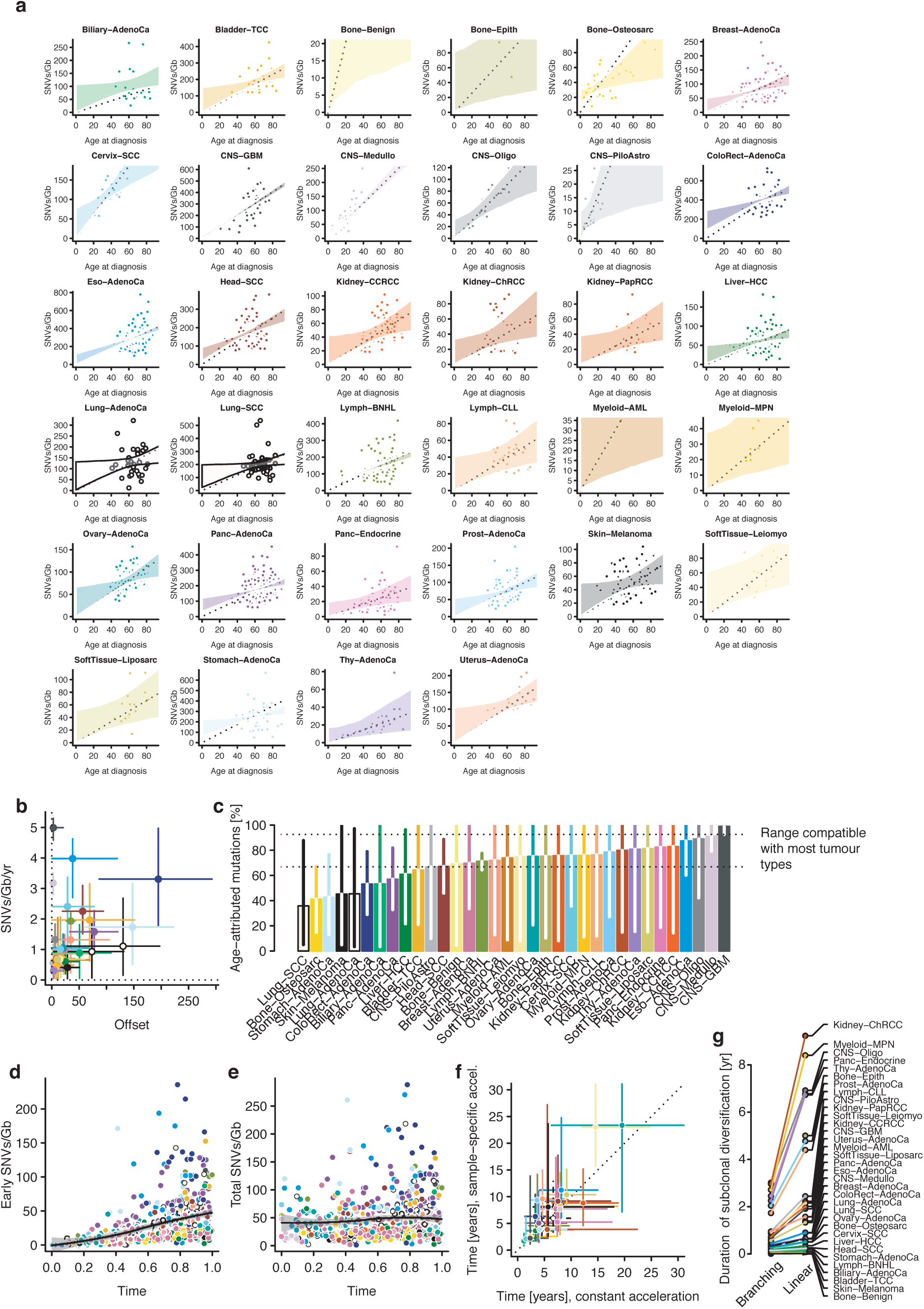
Absolute timing, related to Figure 6. (**a**)CpG>TpG mutation/Gb (scaled to copy number) as a function of age at diagnosis for 33 informative cancer types. The dotted line denotes the median mutations/yr (ie., not offset), while the shading denotes the 95% credible interval of a hierarchical Bayesian linear regression model. (**b**) Maximum a posteriori estimates of rate and offset for 33 cancer types with 95% credible intervals. (**c**) Median fraction of mutations attributed to linear age-dependent accumulation, based on estimates from **b** and the age at diagnosis for each sample. (**d**) Scatter plot of number of early (co-amplified) CpG>TpG mutations (y-axis) as a function of the mutational time estimate of WGD (x-axis). The black line denotes a non-linear loess fit with 95% confidence interval. Colours define the cancer type as in part **a**. (**e**) Total CpG>TpG mutations (y-axis) as a function of the mutation time estimate of WGD (x-axis). Colours and fit as in **d**. (**f**) Inferred median absolute timing patient-specific rate increase, depending on the observed CpG>TpG mutation burden, allowing for a higher (up to 10x) mutation rate increase in cancers with more mutations. (**g)**. Comparison of the median duration of subclonal diversification per cancer type assuming branching and linear phylogenies.

## Author contributions

MG, CJ, IL, SG, PA, DR, DGL, PTS and PVL performed timing of point mutations and copy number gains. SG and MG performed qualitative timing of driver point mutations and analyses of synchronous gains, LJ timed secondary copy number gains. IL, TJM, DR, DGL, DCW and GG performed relative timing of somatic driver events and implemented integrative models. CJ, YR, PVL and QDM performed timing of mutational signatures. MG performed real-time estimation of whole-genome duplication and subclonal diversification. SG assessed mutation rates in relapsed samples. CJ, MG, IL, YR, DR and PVL constructed cancer timelines. MG, CJ, IL, SCD, SG, TJM, YR, PA, JD, PCB, DDB, VM, QDM, PTS, DCW and PVL interpreted the results. SCD, IL, JW, AD, IVG, KeY, GM, MP, SM, ND, KaY, SSe, KH, MT, JD, DGL, DR, JL, MC, SCS, YJ, FM, VM, HZ, WW, QDM, DCW and PVL performed subclonal architecture analysis. SCD, IL, KK, VM, MP, XY, DGL, SSc, RB, MI, MS, DCW and PVL performed copy number analysis. JW, SCD, IL, KH, DGL, KK, DR, DCW, QDM and PVL derived a consensus of copy number analysis results. KaY, MT, AD, SCD, IL, DCW, MG, PVL, QDM and WW derived a consensus of subclonal architecture results. YF and WW contributed to subclonal mutation calls. PTS, DCW and PVL coordinated the study. MG, CJ, PTS, YR, IL, QDM, DCW and PVL wrote the manuscript, which all authors approved.

### Acknowledgements

This work was supported by the Francis Crick Institute, which receives its core funding from Cancer Research UK (FC001202), the UK Medical Research Council (FC001202), and the Wellcome Trust (FC001202). This project was enabled through access to the MRC eMedLab Medical Bioinformatics infrastructure, supported by the Medical Research Council (grant number MR/L016311/1). MT and JD are postdoctoral fellows supported by the European Union’s Horizon 2020 research and innovation program (Marie Sklodowska-Curie Grant Agreement No. 747852-SIOMICS and 703594-DECODE). JD is a postdoctoral fellow of the FWO. FM, GM and KeY would like to acknowledge the support of the University of Cambridge, Cancer Research UK and Hutchison Whampoa Limited. GM, KeY and FM were funded by CRUK core grants C14303/A17197 and A19274. SSe and YJ are supported by NIH R01 CA132897. SM is supported by the Vanier Canada Graduate Scholarship. SCS is supported by the NSERC Discovery Frontiers Project, “The Cancer Genome Collaboratory” and NIH Grant GM108308. HZ is supported by grant NIMH086633 and an endowed Bao-Shan Jing Professorship in Diagnostic Imaging. WW is supported by the U.S. National Cancer Institute (1R01 CA183793 and P30 CA016672). PTS was supported by U24CA210957 and 1U24CA143799. DCW is funded by the Li Ka Shing foundation. PVL is a Winton Group Leader in recognition of the Winton Charitable Foundation’s support towards the establishment of The Francis Crick Institute.

**Members of the PCAWG Evolution and Heterogeneity Working Group**

Stefan C. Dentro^1,2,3,*^, Ignaty Leshchiner^4,*^, Moritz Gerstung^5,*^, Clemency Jolly^1,*^, Kerstin Haase^1,*^, Maxime Tarabichi^1,2,*^, Jeff Wintersinger^6,7,*^, Amit G. Deshwar^6,7,*^, Kaixian Yu^8,*^, Santiago Gonzalez^5,*^, Yulia Rubanova^6,7,*^, Geoff Macintyre^9,*^, David J. Adams^2^, Pavana Anur^10^, Rameen Beroukhim^4,11^, Paul C. Boutros^6,12^, David D. Bowtell^13^, Peter J. Campbell^2^, Shaolong Cao^8^, Elizabeth L. Christie^13,14^, Marek Cmero^14,15^, Yupeng Cun^16^, Kevin J. Dawson^2^, Jonas Demeulemeester^1,17^, Nilgun Donmez^18,19^, Ruben M. Drews^9^, Roland Eils^20,21^, Yu Fan^8^, Matthew Fittall^1^, Dale W. Garsed^13,14^, Gad Getz^4,22,23,24^, Gavin Ha^4^, Marcin Imielinski^25,26^, Lara Jerman^5,27^, Yuan Ji^28,29^, Kortine Kleinheinz^20,21^, Juhee Lee^30^, Henry Lee-Six^2^, Dimitri G. Livitz^4^, Salem Malikic^18,19^, Florian Markowetz^9^, Inigo Martincorena^2^, Thomas J. Mitchell^2,31^, Ville Mustonen^32^, Layla Oesper^33^, Martin Peifer^16^, Myron Peto^10^, Benjamin J. Raphael^34^, Daniel Rosebrock^4^, S. Cenk Sahinalp^19,35^, Adriana Salcedo^12^, Matthias Schlesner^20^, Steven Schumacher^4^, Subhajit Sengupta^28^, Ruian Shi^6^, Seung Jun Shin^8,36^, Lincoln D. Stein^12^, Ignacio Vázquez-García^2,31^, Shankar Vembu^6^, David A. Wheeler^37^, Tsun-Po Yang^16^, Xiaotong Yao^25,26^, Ke Yuan^9,38^, Hongtu Zhu^8^, Wenyi Wang^8,#^, Quaid D. Morris^6,7,#^, Paul T. Spellman^10,#^, David C. Wedge^3,39,#^, Peter Van Loo^1,17,#^

^1^The Francis Crick Institute, London NW1 1AT, United Kingdom; ^2^Wellcome Trust Sanger Institute, Cambridge CB10 1SA, United Kingdom; ^3^Big Data Institute, University of Oxford, Oxford OX3 7LF, United Kingdom; ^4^Broad Institute of MIT and Harvard, Cambridge, MA 02142, USA; ^5^European Molecular Biology Laboratory, European Bioinformatics Institute (EMBL-EBI), Cambridge CB10 1SD, United Kingdom; ^6^University of Toronto, Toronto, ON M5S 3E1, Canada; ^7^Vector Institute, Toronto, ON M5G 1L7, Canada; ^8^The University of Texas MD Anderson Cancer Center, Houston, TX 77030, USA; ^9^Cancer Research UK Cambridge Institute, University of Cambridge, Cambridge CB2 0RE, United Kingdom; ^10^Molecular and Medical Genetics, Oregon Health & Science University, Portland, OR 97231, USA; ^11^Dana-Farber Cancer Institute, Boston, MA 02215, USA; ^12^Ontario Institute for Cancer Research, Toronto, ON M5G 0A3, Canada; ^13^Peter MacCallum Cancer Centre, Melbourne, VIC 3000, Australia; ^14^University of Melbourne, Melbourne, VIC 3010, Australia; ^15^Walter + Eliza Hall Institute, Melbourne, VIC 3000, Australia; ^16^University of Cologne, 50931 Cologne, Germany; ^17^University of Leuven, B-3000 Leuven, Belgium; ^18^Simon Fraser University, Burnaby, BC V5A 1S6, Canada; ^19^Vancouver Prostate Centre, Vancouver, BC V6H 3Z6, Canada; ^20^German Cancer Research Center (DKFZ), 69120 Heidelberg, Germany; ^21^Heidelberg University, 69120 Heidelberg, Germany; ^22^Massachusetts General Hospital Center for Cancer Research, Charlestown, MA 02129, USA; ^23^Massachusetts General Hospital, Department of Pathology, Boston, MA 02114, USA; ^24^Harvard Medical School, Boston, MA 02215, USA; ^25^Weill Cornell Medicine, New York, NY 10065, USA; ^26^New York Genome Center, New York, NY 10013, USA; ^27^University of Ljubljana, 1000 Ljubljana, Slovenia; ^28^NorthShore University HealthSystem, Evanston, IL 60201, USA; ^29^The University of Chicago, Chicago, IL 60637, USA; ^30^University of California Santa Cruz, Santa Cruz, CA 95064, USA; ^31^University of Cambridge, Cambridge CB2 0QQ, United Kingdom; ^32^University of Helsinki, 00014 Helsinki, Finland; ^33^Carleton College, Northfield, MN 55057, USA; ^34^Princeton University, Princeton, NJ 08540, USA; ^35^Indiana University, Bloomington, IN 47405, USA; ^36^Korea University, Seoul, 02481, Republic of Korea; ^37^Human Genome Sequencing Center, Baylor College of Medicine, Houston, TX 77030, USA; ^38^University of Glasgow, Glasgow G12 8RZ, United Kingdom; ^39^Oxford NIHR Biomedical Research Centre, Oxford OX4 2PG, United Kingdom.

^*^: These authors contributed equally

^#^: These authors jointly directed the work

